# Gene activation by a CRISPR-assistant *trans* enhancer

**DOI:** 10.1101/517920

**Authors:** Xin Hui Xu, Wei Dai, Dan Yang Wang, Jian Wu, Jin Ke Wang

**Affiliations:** State Key Laboratory of Bioelectronics, Southeast University, Nanjing 210096, China

**Keywords:** CRISPR/dCas9, CMV enhancer, trans enhancer, gene activation

## Abstract

Gene activation is essential to the basic biological research and biomedicine. Therefore, various gene activators such as activation domain-ZNF, TALE and CRISPR proteins have been developed for this end, in which the CRISPR protein dead Cas9 (dCas9) is now most widely used. However, the current gene activators are still limited by their inefficient gene activation activity. In this study, we developed a new strategy, CRISPR-assistant *trans* enhancer, for activating gene expression in high efficiency by combining dCas9-VP64/sgRNA with a widely used strong enhancer, the CMV enhancer. In this strategy, a *trans* CMV enhancer DNA was recruited to target gene by dCas9-VP64/sgRNA via annealing between 3’ end of sgRNA and CMV enhancer. The *trans* enhancer activates gene transcription as the natural looped *cis* enhancer. The *trans* enhancer could activate both exogenous reporter gene and variant endogenous genes in various cells, with much higher activation efficiency than the current dCas9 activators.

## Introduction

Artificial activation of gene expression has important role in the basic biological research and biomedical application. For example, gene function is often explored by artificially activating its expression in cells or *in vivo* in basic research. In biomedicine, cells are often needed to be reprogrammed into induced stem (iPS) cells or other differentiated cells by activating endogenous genes. In medicine, cancers can be treated by inhibiting various kinases, activating genes enhancing immunity (**Sagiv-Barfi *et al*. 2018**), apoptosis, and differentiation (**McClellan *et al*. 2015**). Therefore, various artificial gene activators have been developed. For instance, the activation domain-fused zinc finger (ZNF), transcription activator-like effector (TALE), and clustered regularly interspaced short palindromic repeats (CRISPR) associated proteins have been developed for gene activation. In these proteins and complexes, the CRISPR-associated (Cas) proteins are now most widely used due to its simplicity (**Gao *et al*. 2014b**).

CRISPR is originally an immune system of bacteria to destroy the invaded microphage DNAs by enzymatically digestion. The system has already been developed into a high efficient gene editing tool (**Doudna and Charpentier 2014; Mali *et al*. 2013c**). In addition, the system has also been developed as a new kind of gene regulation tool. For example, the dead Cas9 (dCas9) and its associated single guide RNA (sgRNA) have been most widely used to regulate gene expression in recent years (**Dominguez *et al*. 2015; Hilton *et al*. 2015; Jinek *et al*. 2012; Kiani *et al*. 2015; Mali *et al*. 2013b; Radzisheuskaya *et al*. 2016; Wang *et al*. 2016**). For this end, both dCas9 and sgRNA have been widely engineered for activating or repressing gene expression. For instance, the dCas9 protein was fused with various gene activation or repression domains, such as VP48 (**Cheng *et al*. 2013**), VP160 (**Perrin *et al*. 2017**), VP64 (**Maeder *et al*. 2013; Perez-Pinera *et al*. 2013**), VPR (VP64-p65-Rta) (**Chavez *et al*. 2015**), and KRAB (Zheng *et al*. 2018). Additionally, the dCas9 protein was also fused with other functional domains with transcriptional regulatory functions, such as p300(**Hilton *et al*., 2015**), LSD1(**Kearns *et al*. 2015**), Dnmt3a (**Liu *et al*. 2016a; Saunderson *et al*. 2017**), and Tet1(**Choudhury *et al*. 2016; Liu *et al*., 2016a**). Based on these domains, more elaborate activators have been developed for more potent activation of target genes in mammalian cells, such as SunTag (dcas9-GCN4/sgRNA plus scFV-VP64) (Tanenbaum *et al*. 2014), and SPH (dCas9-GCN4/sgRNA plus scFV-p65-HSF1) (**Zhou *et al*. 2018**). More complex, some inducible dCas9 systems have been developed to control activity of dCas9 activators in cells, such as light-activated CRISPR/Cas9 effector (**Nihongaki *et al*. 2015; Polstein and Gersbach 2015**), hybrid drug inducible CRISPR/Cas9 technology (HIT) (**Lu *et al*. 2018**), and CRISPR activator gated by human antibody-based chemically induced dimerizers (AbCIDs) (**Liu *et al*. 2018**). A trouble of these typical dCas9 fusion proteins in their *in vivo* applications is that they are difficult to be accommodated by adeno-associated virus (AAV), a most suitable type of non-integrating gene therapy vector, due to its limited packaging capacity.

Besides dCas9 engineering, sgRNA has also been engineered to develop new dCas9-based activators. Compared with dCas9 engineering, sgRNA is more simple, flexible and efficient to redesign. Moreover, the engineered sgRNA is more helpful for the *in vivo* application of dCas9-based activators due to its limited length for virus packaging. The most widely used engineered sgRNA is the sgRNA fused with MS2 loops at the 3’ end (sgRNA-MS aptamer), which can be bound by dimerized MS2 bacteriophage coat proteins fused with transcription-activating domains VP64-HSF1 (MS2-VP64-HSF1, MPH)(**Konermann *et al*. 2015; Liao *et al*. 2017**). The system is now well known as the synergistic activation mediator (SAM) system. Similarly, another engineered sgRNA-based dCas9 activator, named as the Casilio system, was developed, which consists of the dCas9 protein, an sgRNA appended with one or more binding site(s) of RNA-binding protein Pumilio/FBF (PUF) (sgRNA-PBS), and an effector (such as VP64 and p65-HSF1) fused with a PUF domain (PUF fusion) (**Cheng *et al*. 2016**). In the same way, by extending sgRNAs to include effector protein recruitment sites, the modular scaffold RNAs that encode both target locus and regulatory action were constructed (**Zalatan *et al*. 2015**). For these recruitment RNA modules, the well-characterized viral RNA sequences MS2, PP7, and com, which are recognized by the MCP, PCP, and Com RNA-binding proteins, respectively, were used (**Zalatan *et al*., 2015**). The transcriptional activation domain VP64 was fused to each of the corresponding RNA-binding proteins (**Zalatan *et al*., 2015**). By extending sgRNAs to include modified riboswitches that recognize specific signals, a CRISPR-Cas9-based ‘signal conductors’ that regulate transcription of endogenous genes in response to external or internal signals of interest (such as drug) was created (**Liu *et al*. 2016b**). Clearly, these chimeric sgRNAs limited by their long complexed RNA aptamers and the cognate RNA-binding fusion proteins.

Although variant dCas9-based activators have been developed (**Chen and Qi 2017**), the current dCas9-based transcriptional activators are still inefficient at endogenous gene activation and reprogramming (**Gao *et al*. 2014a**). By a systematic comparing of their relative potency and effectiveness across various cell types and species (human, mouse, and fly) (**Chavez *et al*. 2016**), it was found that the majority of second-generation activators showed improved levels of activation as compared to those of dCas9-VP64, in which three activators including VPR, SAM, and Suntag were most potent. Across a range of target genes and cellular environments, the VPR, SAM, and Suntag systems are consistently superior to the previous VP64 standard. Moreover, VPR, SAM, and Suntag generally fall within an order of magnitude of each other with regard to fold increase in gene expression. More importantly, attempts to build an improved chimeric activator by fusing elements from VPR, SAM, and Suntag were unsuccessful (**Chavez *et al*., 2016**). Therefore, future efforts to improve dCas9-based activators may benefit from exploring other unique architectures or novel activation domains.

Almost three decades ago, the human cytomegalovirus (CMV) enhancer/promoter (referred to as CMV enhancer hereafter) was found as a natural mammalian promoter with high transcriptional activity (**Boshart *et al*. 1985**). The late studies gradually found that the CMV enhancer is the known strongest promoter in various mammalian cells (**Boshart *et al*., 1985; Foecking and Hofstetter 1986; Ho *et al*. 2015; Kim *et al*. 1990**). Therefore, this enhancer has been widely used to drive the ectopic expression of various genes in wide range of mammalian cells. For example, the CMV enhancer is also used to drive the ectopic expressions of exogenous genes in broad tissues in transgenic animals (**Furth *et al*. 1991; Schmidt *et al*. 1990**), protein production by gene engineering, and human gene therapy. We have recently further improved the transcriptional activity of the CMV enhancer by changing the natural NF-κB binding sites in this enhancer into artificially selected NF-κB binding sequences with high binding affinity (**Wang *et al*. 2018**). Therefore, we conceived that an unique architecture may be constructed to further improve dCas9-based activators by using the CMV enhancer.

In this study, mimicking the natural enhancer activating gene expression by a loop structure (**Carter *et al*. 2002; Tolhuis *et al*. 2002**), we developed a new dCas9-based activator by combining dCas9/sgRNA with CMV enhancer. The 3’ end of sgRNA was redesigned to add a short capture sequence in complementary with a stick-end of a double-stranded CMV enhancer. The CMV enhancer is anchored to the promoter region of target gene by dCas9/sgRNA. The dCas9/sgRNA-recruited CMV enhancer thus functions like a natural looped *cis* enhancer in *trans* form. We found that the new activator could efficiently activate multiple genes in 6 kinds of cells including 293T, HepG2, PANC-1, HeLa, A549, and HT29. Moreover, the new activator activated expression of transcription factors *HNF4α* in HepG2 cells and *E47* in PANC-1 cells could lead to differentiation of these cancer cells.

## Results

### Conception of gene activation by a CRISPR-assistant *trans* enhancer

The principle of activating gene expression by a CRISPR-assistant *trans* enhancer is schematically illustrated in **Figure 1a**. We modified sgRNA by adding a capture sequence to the 3’ end of normal sgRNA sequence, which produces a 3’ end-extended sgRNA. Because the newly designed sgRNA will be used to capture a *trans* enhancer DNA, we named it as capture sgRNA (csgRNA). Correspondingly, a linear double-stranded CMV enhancer sequence with a free single-strand 3’ end was designed; we named the CMV fragment as stick CMV (sCMV), which can interact with csgRNA by annealing to the 3’ capture sequence of csgRNA via its stick end. In our conception, with these csgRNA and sCMV designing, when the csgRNA directs the dCas9 protein to the target site that locates in the promoter region of interested gene, the sCMV will be captured to the gDNA-bound dCas9/csgRNA. This interaction process will anchor a sCMV to the promoter region of an interested gene. The anchored sCMV may thus activate the transcription of an interested gene like natural looped *cis* enhancer. Because the dCas9/csgRNA-anchored sCMV functions like transcription factors in *trans*, it can be regarded as *trans* enhancer, in contrast with natural enhancer that functions in *cis*.

**Figure 1.**
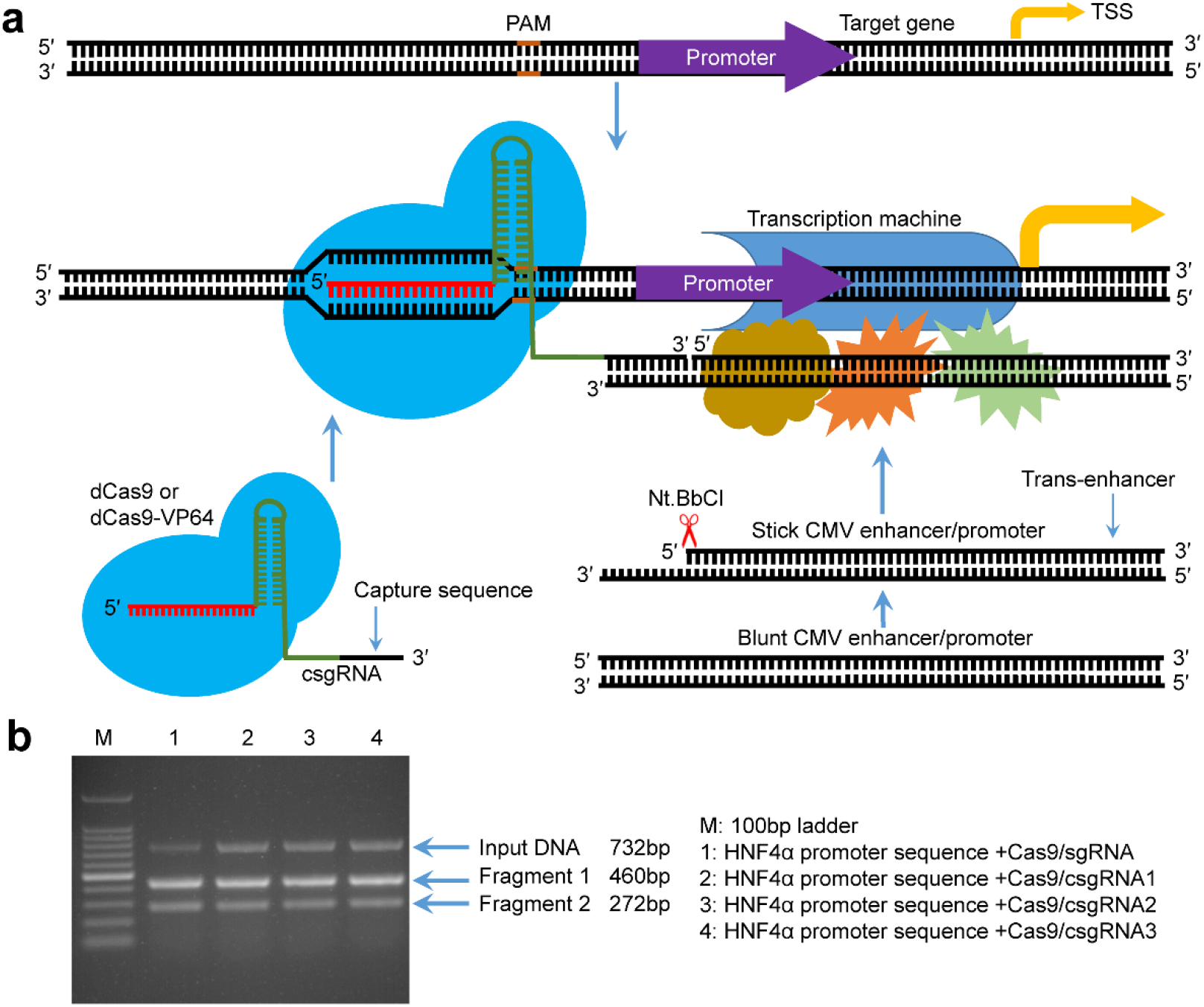
Principle of gene expression activation by the CRISPR-assistant *trans* enhancer and evaluation of designed csgRNAs. a. Schematic illustration of the principle of gene expression activation by the CRISPR-assistant *trans* enhancer. A capture sequence was added at the 3’ end of sgRNA, which is used to capture a *trans* CMV enhancer with a single-stranded overhang that can anneal with the capture sequence of sgRNA. The captured *trans* CMV enhancer may function like the natural looped *cis* enhancer to activate transcription of interested gene, including exogenous and endogenous genes. b. *In vitro* target DNA cutting by the Cas9-csgRNA complex. A 732-bp DNA fragment amplified from the HNF4α promoter region were respectively cut by the Cas9/csgRNA and Cas9/sgRNA complexes.

### Effect of capture sequence on the function of sgRNA

An extended capture sequence was added to the sgRNA scaffold for annealing with the stick end of sCMV enhancer fragment. To find whether the added capture sequence affects the function of sgRNA, we synthesized a normal sgRNA and three csgRNAs targeting to a same site of *HNF4α* promoter by *in vitro* transcription. Three csgRNAs had different capture sequences. We used these sgRNAs to associate with the Cas9 endonuclease to cut a 732-bp *HNF4α* promoter fragment. The results indicate that the target DNA could be digested by all sgRNAs, including three csgRNAs with variant capture sequences (**Figure 1b**). This indicates that the capture sequence exerts no effect on the sgRNA function.

### Activation of an exogenous reporter gene by *trans* enhancer

To find whether the CRISPR-assistant *trans* enhancer is able to activate gene expression, we constructed a reporter construct of *HNF4α* promoter, pEZX-HP-ZsGreen. We then transfected the 293T cells with sCMV, dCas9, and csgRNA2. We found that the system activated the expression of reporter gene ZsGreen (**Figure 2**; dCas9/csgRNA2-sCMV). The activation level is similar to that of dCas9-VP64/sgRNA. Because we used a dCas9 that has no fused transcriptional activation domain like VP64 in the trial transfection, this indicates that the sCMV can not only interact with the dCas9/csgRNA in cell as we conceived, but also function as a transcription factor to activate gene expression as we expected. However, the activation level of dCas9/csgRNA & sCMV system is far below the activation strength of *cis* CMV enhancer.

To improve the transcriptional activation performance of dCas9/csgRNA & sCMV system, we then transfected the 293T cells with a dCas9-VP64/csgRNA2 & sCMV system. As a result, we found that the system greatly activated the expression of reporter gene (**Figure 2**; dCas9-VP64/csgRNA2-sCMV). As a control, the transfection with dCas9-VP64/csgRNA2 & blunt CMV (bCMV) system obtained an activation level similar to that of dCas9-VP64/sgRNA (**Figure 2**), indicating that the sCMV genuinely contributed to the gene expression activation. More importantly, we found that the VP64 fused to the dCas9 protein could significantly improve the function of sCMV, possibly by addition or synergy. We then transfected the 293T cells with other two csgRNAs, csgRNA1 and csgRNA3, together with dCas9-VP64 & sCMV. We found that these two csgRNAs also obtained high level activation (**Figure 2**), despite inferior to csgRNA2. The transcriptional activation function of *trans* sCMV was also confirmed by another control transfection, dCas9/csgRNA2 & bCMV, which showed no transcriptional activation (**Figure 2**).

**Figure 2.**
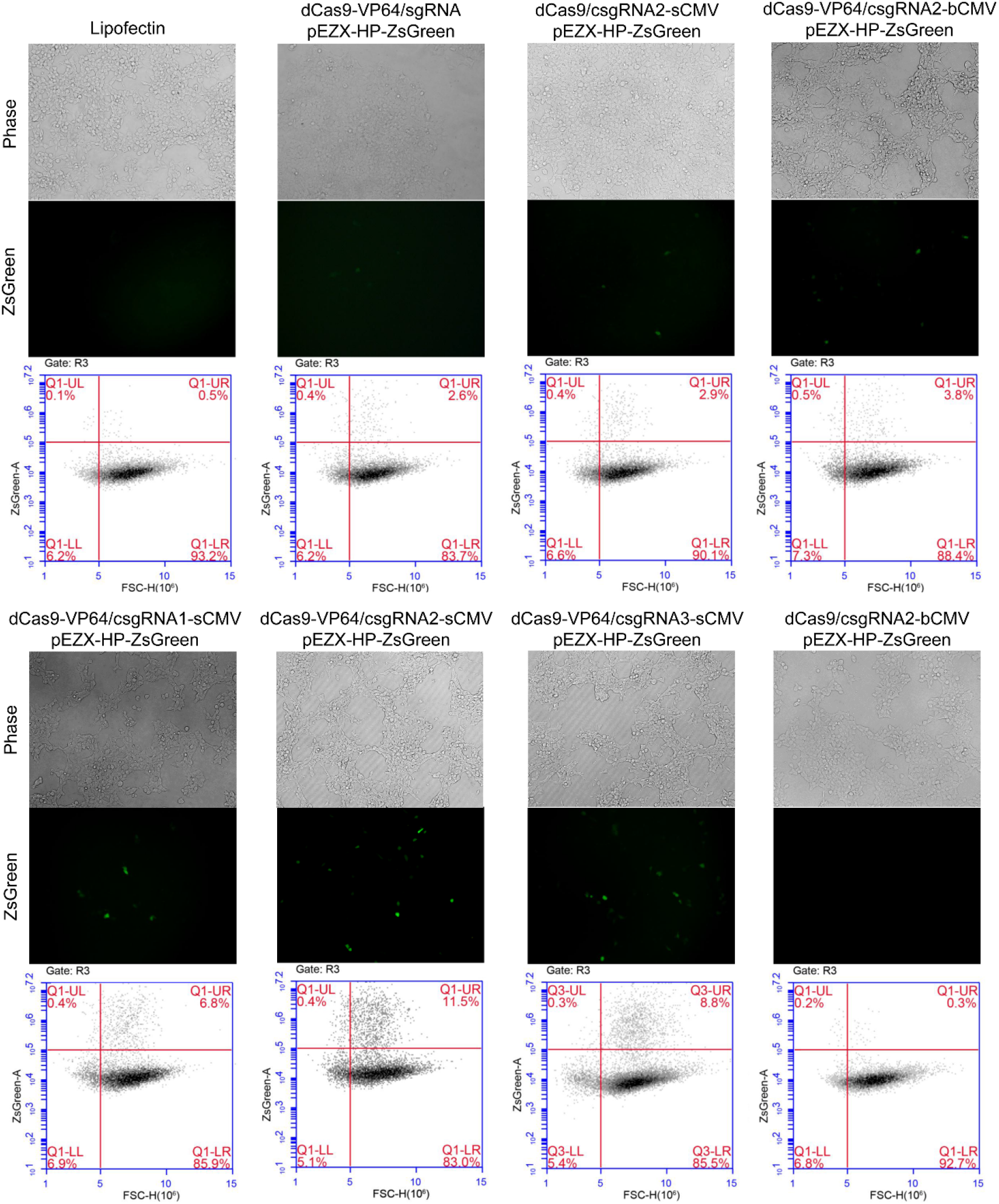
Activation of an exogenous reporter gene ZsGreen under the control of a *HNF4α* promoter by the CRISPR-assistant *trans* enhancer in 293T cells. Cells were transfected by various vectors. Cells were photographed with a fluorescent microscope and their florescence was analyzed by flow cytometry. In order to reveal the gene activation activity, variant transfections as controls were simultaneously performed. The reporter gene activation efficiency was indicated by the percentage of cells with green fluorescence over the threshold (cells in Q1-UR quadrant).

To confirm the function of *trans* enhancer, we also performed more negative control transfections in 293T cells, including the sole dCas9-VP64, sgRNA, and pEZX-HP-ZsGreen, and dCas9-VP64 plus pEZX-HP-ZsGreen. These transfections all didn’t activate the reporter gene expression (**Supplementary Figure 1**). These data verify that the *trans* enhancer sCMV can play transcriptional activation role by combination with dCas9/csgRNA. Moreover, the typical transcription activation domain VP64 fused to dCas9 can further improve the transcriptional activation performance of *trans* enhancer sCMV. Therefore, we adopted the dCas9-VP64/csgRNA2 & sCMV system to perform subsequent more characterizations of CRISPR-assistant *trans* enhancer.

The transfection of 293T cells revealed that the dCas9-VP64 & csgRNA2 & sCMV system has the highest transcriptional activation capability. To verify the versatility of the *trans* enhancer system in variant cells, we transfected 6 cell lines, including HepG2 (**Figure 3**), A549 (**Supplementary Figure 2**), HeLa (**Supplementary Figure 3**), SKOV3 (**Supplementary Figure 4**), PANC-1 (**Supplementary Figure 5**), and HT29 (**Supplementary Figure 6**), with the dCas9-VP64 & csgRNA2 & sCMV system and reporter construct. As controls, all cells were also co-transfected by dCas9-VP64 & sgRNA and dCas9 & csgRNA2 & sCMV in combination with reporter construct. The results were similar to those of 293T cells. The dCas9-VP64 & csgRNA2 & sCMV showed the highest transcriptional activation efficiency in all cell lines (**Figure 3, Supplementary Figures 2–6**). It should be noted that the dCas9 & csgRNA2 & sCMV system always showed higher transcriptional activation efficiency than the dCas9-VP64 & sgRNA in all cell lines (**Figure 3, Supplementary Figures 2–6**), which confirms the transcriptional activation capability of *trans* enhancer because the dCas9 & csgRNA2 & sCMV system contains no transactivation domain such as VP64. The highest performance of dCas9-VP64 & csgRNA2 & sCMV system in all transfected cell lines demonstrates that the *trans* sCMV has a synergistic effect with the dCas9-fused transactivation domain VP64 in activating gene transcription. These observations were further supported by the quantified fluorescence intensity (MFI) of cells of two biological replicate transfections (**Figure 4a**).

**Figure 3.**
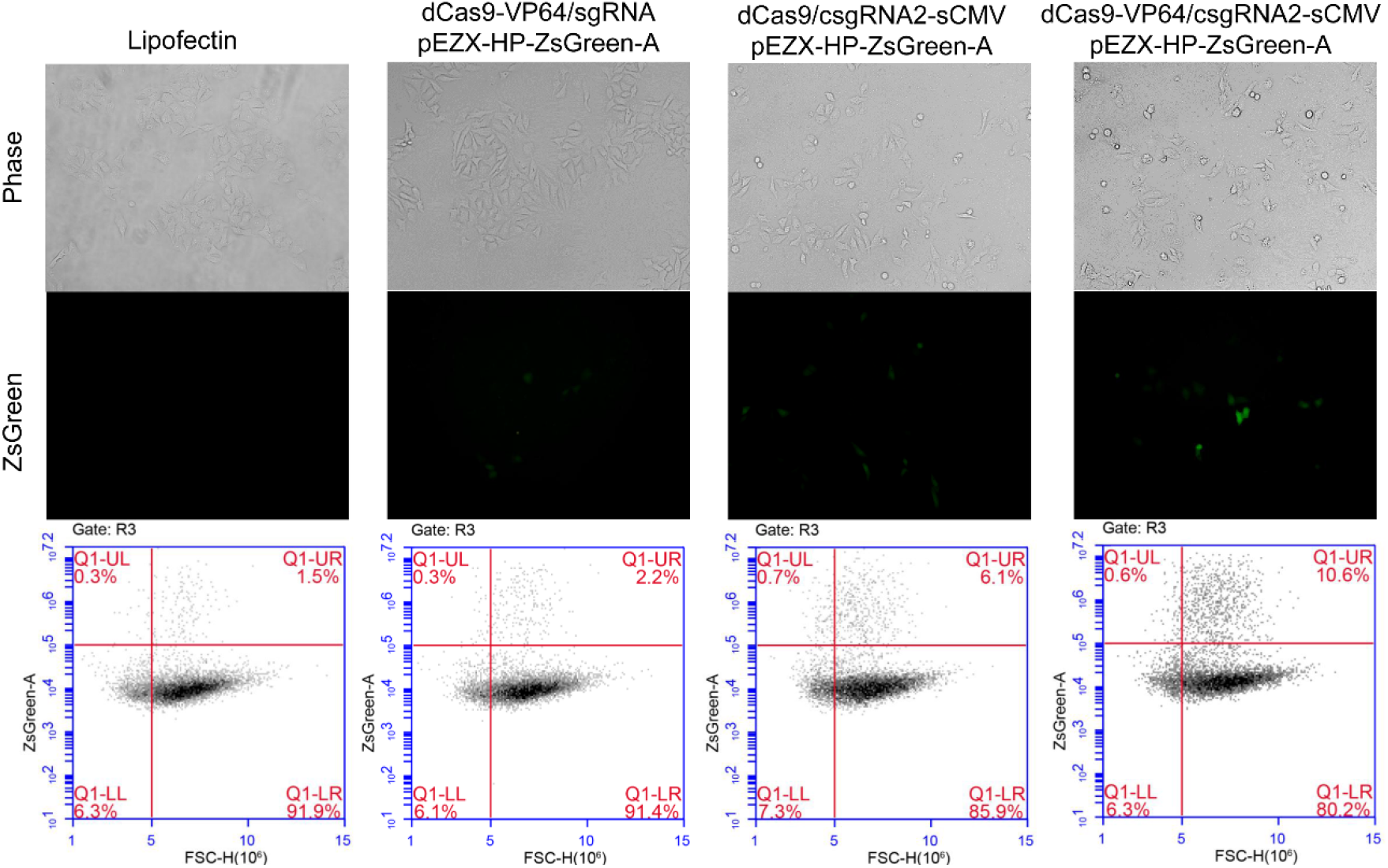
Activation of an exogenous reporter gene ZsGreen under the control of a *HNF4α* promoter by the CRISPR-assistant *trans* enhancer in the HepG2 cells. Cells were transfected with three transcriptional activation systems, including dCas9-VP64 & sgRNA, dCas9 & csgRNA & sCMV, and dCas9-VP64 & csgRNA2 & sCMV, together with a reporter construct (pEZX-HP-ZsGreen-A), respectively. Cells were photographed with a fluorescent microscope and their florescence intensity was analyzed by flow cytometry. The reporter gene activation efficiency was indicated by the percentage of cells with green fluorescence over the threshold (cells in Q1-UR quadrant).

**Figure 4.**
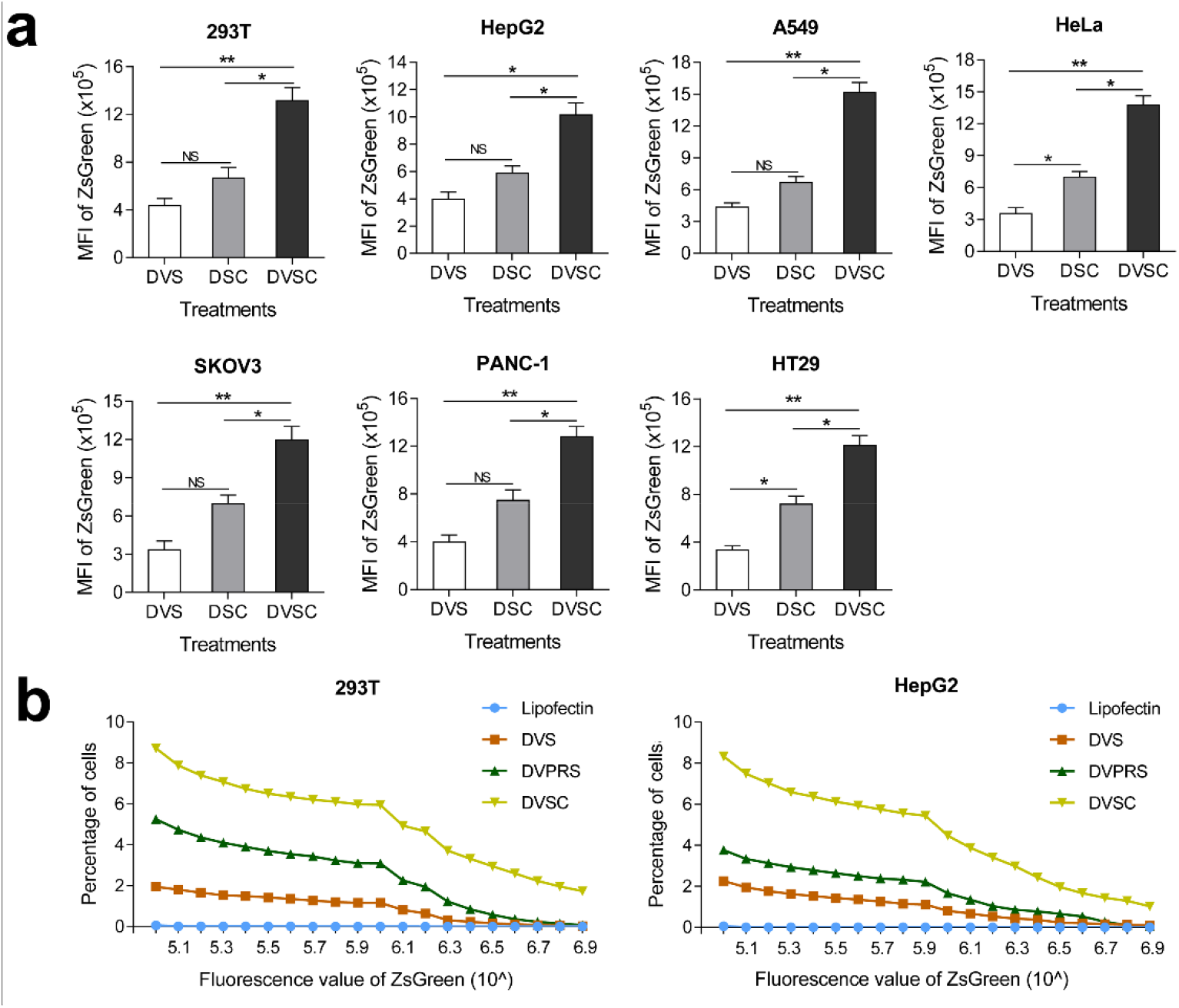
Activation of an exogenous reporter gene ZsGreen under the control of a *HNF4α* promoter by the CRISPR-assistant *trans* enhancer in multiple cells. a. Transcriptional activation of reporter gene ZsGreen in various cells transfected by different vectors. The florescence intensity of cells were analyzed by flow cytometry and showed as the mean fluorescence intensity (MFI). Co-transfections: DVS, dCas-VP64 & sgRNA; DSC, dCas9 & csgRNA & sCMV; DVSC, dCas-VP64 & sgRNA & sCMV. b. Comparison of *trans* enhancer with VPR. Cells were transfected with three different transcriptional activation systems to activate reporter gene ZsGreen. The florescence intensity of cells was analyzed by flow cytometry and the percentage of cells with particular fluorescence intensity was counted. Transfections: Lipo, lipofection; DVS, dCas-VP64 & sgRNA; DVPRS; dCas9-VPR & csgRNA; DVSC, dCas9-VP64 & csgRNA & sCMV.

### Comparison of *trans* CMV enhancer with VPR

The VPR transcription-activating domain has been shown with higher transcriptional activation ability than VP64. Therefore, we next compared the *trans* enhancer with this strong transcription activation domain. We transfected the 293T and HepG2 cells with dCas9-VP64/csgRNA, dCas9-VPR/csgRNA, and dCas9-VP64/csgRNA & sCMV, in combination with ZsGreen reporter construct, respectively. The results reveal that dCas9-VPR/csgRNA had better transcriptional activation efficiency than dCas9-VP64/csgRNA as reported by many previous studies (**Figure 4b**). However, the dCas9-VP64/csgRNA & sCMV showed far higher transcriptional activation efficiency than dCas9-VPR/csgRNA (**Figure 4b**). Moreover, we found that the former activated the ZsGreen expression in more cells than the latter at any fluorescence intensity threshold. This means that the *trans* enhancer sCMV not only make more cells produce fluorescence, but also make more cells produce higher fluorescence than dCas9-VPR/sgRNA. These data further demonstrate the great transcriptional activation ability of CRISPR-assistant *trans* enhancer.

### Activation of endogenous genes by *trans* enhancer

To investigate the transcriptional activation of endogenous genes by *trans* enhancer, we chose 10 genes as activating targets, including *HNF4α, E47* (*TCF3*), *ASCL1, Ngn2, Oct4, Nanog, TNFAIP3, CASP9, CSF3*, and *Sox2*. We designed one promoter-targeting sgRNA for each of these genes. We then transfected 7 different cell lines with the dCas9-VP64/csgRNA2 & sCMV. At the same time, all cells were simultaneously transfected with dCas9-VP64/sgRNA and dCas9/csgRNA & sCMV as controls. After the gene transcription was detected by qPCR, the expression fold of target genes in the cells transfected by dCas9/csgRNA2 & sCMV relative to those only transfected by Lipofectamine 2000 was calculated. By comparing the transcriptional activation effects of these 10 genes in different cell lines, we found that the dCas9-VP64/csgRNA2 & sCMV always activated the highest transcription of all genes in all cells (**Figure 5**). As in exogenous gene activation, the dCas9/csgRNA & sCMV could better activate target genes than dCas9-VP64/sgRNA in all cells. Interestingly, we found that the expression of 6 genes including *HNF4α, TCF3, ASCL1, Ngn2, TNFAIP3*, and *CSF3* were always highly activated by three systems in all cells. However, the expression of other 4 genes including *Oct4, Sox2, Nanog*, and *CASP9*, was just less activated. These results demonstrate that the CRISPR-assistant *trans* enhancer can activate the expressions of variant endogenous genes in various cells. These results also indicate that the *trans* sCMV has a synergistic effect with the dCas9-fused transactivation domain VP64 in activating gene transcription.

**Figure 5.**
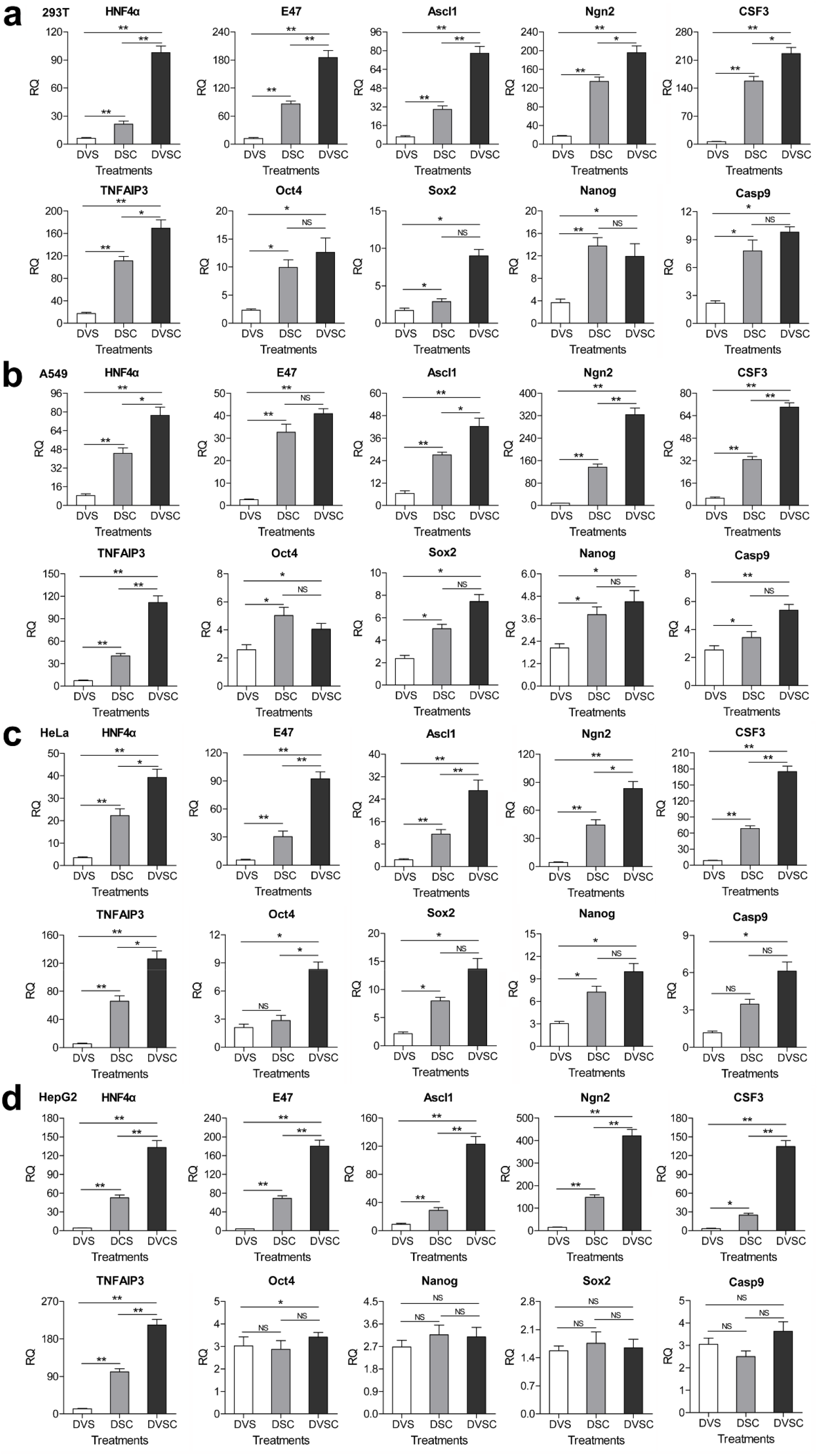
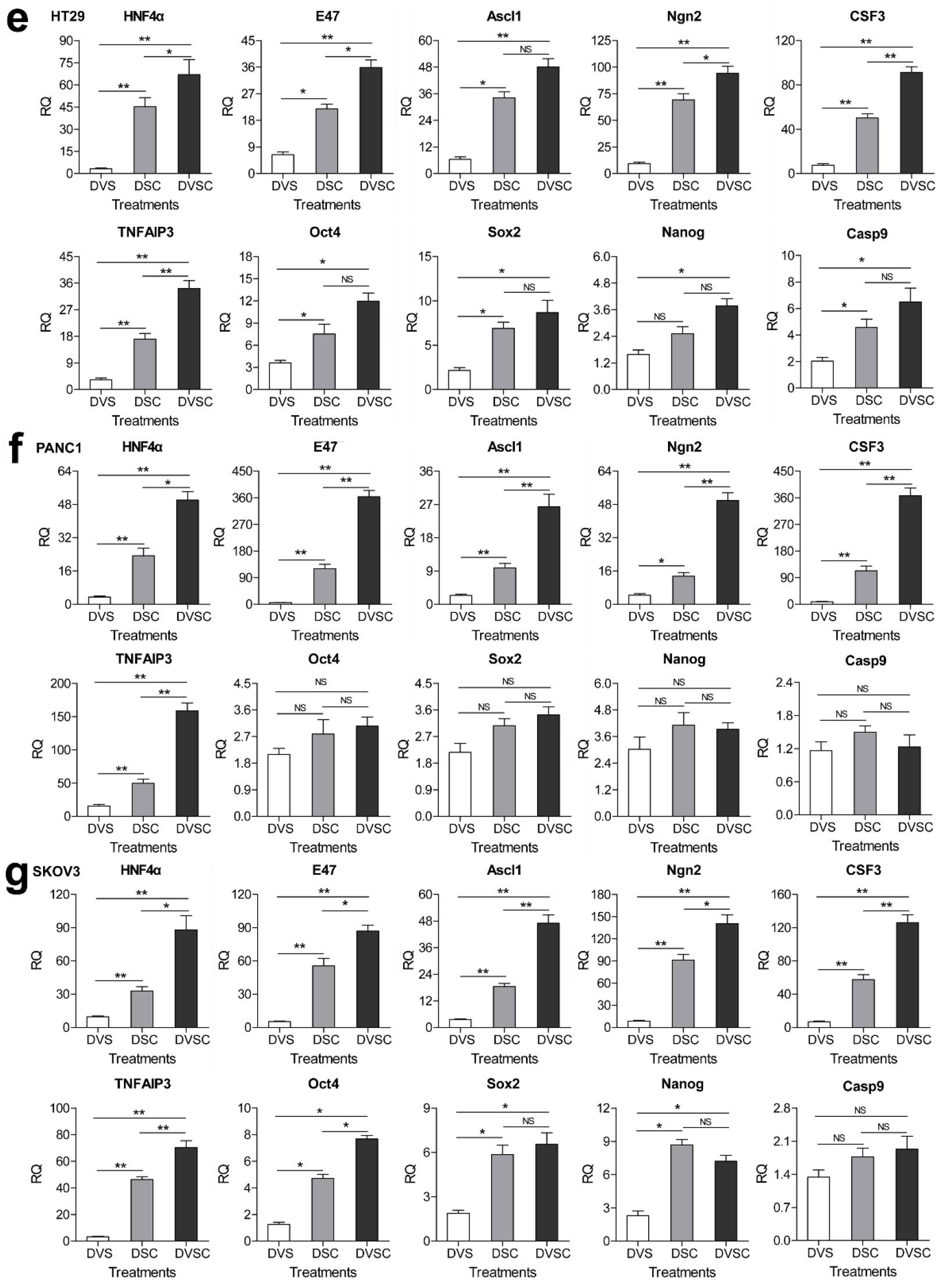
Transcriptional activation of endogenous genes by the CRISPR-assistant *trans* enhancer. Seven different cell lines were transfected with three different transcriptional activation systems to activate the expression of 10 endogenous genes. The gene transcription was detected by qPCR and the expression level was showed as the relative RNA expression fold to house-keeping gene β actin. Three biological replicates were performed for the transfection of each cell line. Data were shown as mean ± SD, n=3. The statistical difference was analyzed by Student’s t-test. *, *P* < 0.05; **, *P* < 0.01; NS, no significant statistical difference. Co-transfection: DVS, dCas9-VP64 & sgRNA; DSC, dCas9 & csgRNA2 & sCMV; DVSC, dCas9-VP64 & csgRNA2 & sCMV.

### Activation of tumor cell differentiation genes by *trans* enhancer

Finally, we explored whether the *trans* enhancer can be used to activate critical genes related to tumor differentiation therapy. We selected two transcription factors that were reported to induce tumor cell differentiation, *HNF4α* and *E47*. The former has been reported to induce the differentiation of cancer cell HepG2, and the latter has been reported to induce the differentiation of cancer cell PANC-1.

We firstly transfected the HepG2 and PANC-1 cells with the dCas9/csgRNA & sCMV system that targets to a single site in the promoter region of *HNF4α* and *E47* genes, respectively. The qPCR detection of gene expression indicates that the two genes were highly activated in the transfected cancer cells (**Figure 5**; see HepG2 and PANC-1 cells). Moreover, with the activation of *HNF4α* in HepG2 cells, the expression of *CD133* and *CD90* was down-regulated (**Figure 6a**). In contrast, the expression of *p21* was highly up-regulated. The expression of some typical genes that are involved in the establishment or maintenance of pluripotency, including *Oct3/4, Sox2, Nanog, c-Myc, LIN28*, and *Klf4*, was also down-regulated (**Figure 6a**). On the contrary, the expression of multiple genes related to healthy liver function, including *GS, BR, ALDOB, CYP1a2, PEPCK, APOCIII, G-6-P*, and *HPD*, was highly up-regulated (**Figure 6b**).

**Figure 6.**
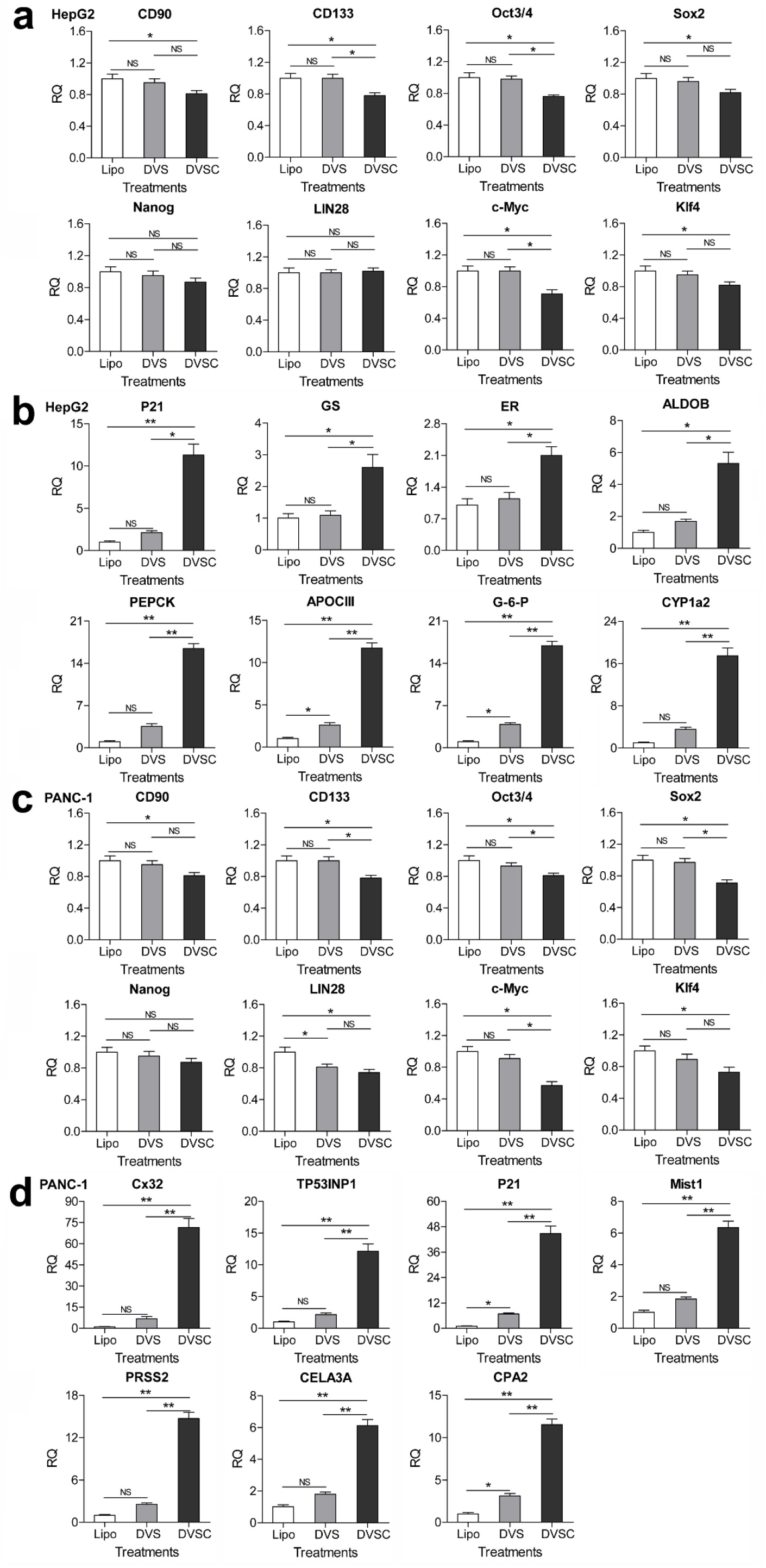
Changes of gene expression and physiological phenotypes in the *HNF4α*-activated HepG2 cells and *E47*-activated PANC-1 cells. a and b. Change of gene expression in the *HNF4α*-activated HepG2 cells. c and d. Change of gene expression in the *E47*-activated PANC-1 cells. The gene transcription was detected by qPCR and the expression level was showed as the relative RNA expression fold to house-keeping gene β actin. Three biological replicates were performed for the transfection of each cell line. Data were shown as mean ± SD, n=3. The statistical difference was analyzed by Student’s t-test. *, *P* < 0.05; **, *P* < 0.01; NS, no significant statistical difference. Transfection: Lipo, lipofectin; DVS, dCas9-VP64 & sgRNA; DVSC, dCas9-VP64 & csgRNA2 & sCMV.

Similarly, when the E47 gene was activated in the PANC-1 cell, the expression of *CD133* and *CD90* was down-regulated (**Figure 6c**). In contrast, the expression of *p21* and *TP53INP1* was highly up-regulated. The expression of some typical genes that are involved in the establishment or maintenance of pluripotency, including *Oct3/4, Sox2, Nanog, c-Myc, LIN28*, and *Klf4*, was down-regulated (**Figure 6c**), while the expression of several genes related to healthy pancreas function, including *MIST1, PRSS2, CELA3A*, and *CPA2*, was significantly up-regulated (**Figure 6d**). The *p21* and *TP53INP1* was required by E47-induced cell cycle arrest (**Figure 6d**). The *MIST1* normally regulated the acinar maturation pathways. The gene *PRSS2, CELA3A*, and *CPA2* code the digestive enzyme trypsin, elastase 3, and carboxypeptidase A2, respectively.

Besides detecting gene expression, we also detected the physiological phenotypes of transfected cells, including proliferation, migration, and invasion. We characterized the cell proliferation by detecting cell cycles. The results indicate that the *HNF4α* and *E47* activation by *trans* enhancer induced significant cell growth arrest (increased G_0_/G_1_ cells) in HepG2 and PANC-1 cells, respectively (**Figure 7a**). The wound-healing assay reveals that the migration ability of two cell lines was significantly decreased by the transfection of *trans* enhancer (**Figure 7b, Supplementary Figure 7**). The transwell assay also revealed that the *trans* enhancer transfection resulted in significant decrease of invasion ability of two cell lines (**Supplementary Figure 8**). These physiological changes of the transfected cells are in agreement with the gene expression changes detected above.

**Figure 7.**
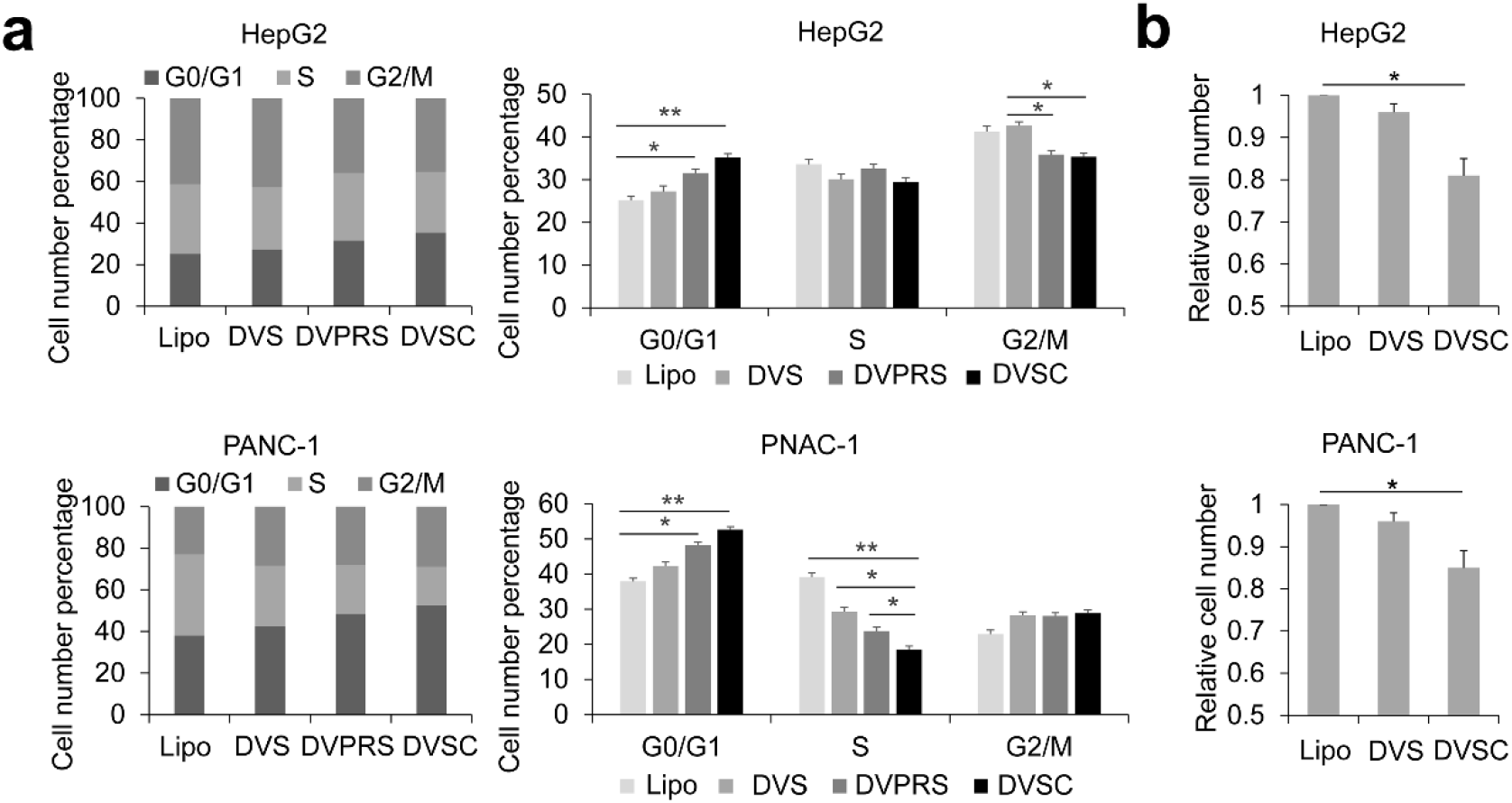
Change of cell cycle and migration in the *HNF4α*-activated HepG2 cells and *E47*-activated PANC-1 cells. a. Change of cell cycle in the *HNF4α*-activated HepG2 cells and *E47*-activated PANC-1 cells. b. Change of cell migration in the *HNF4α*-activated HepG2 cells and *E47*-activated PANC-1 cells. Three biological replicates were performed for the transfection of each cell line. Data were shown as mean ± SD, n=3. The statistical difference was analyzed by Student’s t-test. *, *P* < 0.05; **, *P* < 0.01; NS, no significant statistical difference. Transfection: Lipo, lipofectin; DVS, dCas9-VP64 & sgRNA; DVPRS, dCas9-VPR & csgRNA2; DVSC, dCas9-VP64 & csgRNA2 & CMV.

### Activation of endogenous gene by *trans* enhancer consisting of VPR and CMV

Finally, we expected to known whether the combination of dCas9-VPR/csgRNA and sCMV could obtain higher transcriptional activation than the combination of dCas9-VP64/csgRNA and sCMV. We thus transfected 293T cell with dCas9-VP64 and dCas9-VPR together with csgRNA2 and sCMV to activate the endogenous gene *HNF4α*. The results indicated that dCas9-VPR/csgRNA & sCMV always obtained significant lower activation of *HNF4α* expression than dCas9-VP64/csgRNA & sCMV (**Supplementary Figure 9**).

## Discussion

As a typical application, the dCas9-based transcriptional activators have recently been used to reprogram cells *in vitro* and *in vivo* for biomedical applications by activating endogenous genes. For example, the fibroblasts can be reprogramed to pluripotency (iPS cell) by activating endogenous Oct4 and Sox2 with dCas9-SunTag-VP64 (**Liu *et al*., 2018**). The mouse embryonic fibroblasts can be converted to induced neuronal cells by activating endogenous *Brn2, Ascl1*, and*Myt1l* genes with^VP64^dCas9^VP64^ (**Black *et al*. 2016**). The in vivo target genes can be activated by MPH to ameliorate disease phenotypes in mouse models of type I diabetes, acute kidney injury, and muscular dystrophy (**Liao *et al*., 2017**). The brain astrocytes can be directly and efficiently converted into functional neurons *in vivo* by activating *Ascl1, Neurog2* and *Neurod1* genes with SPH (**Zhou *et al*., 2018**). These studies makes CRISPR therapies the grade not the cut (**Burgess 2018**). However, dCas9-based transcriptional activators are still inefficient at endogenous gene activation and reprogramming (**Gao *et al*., 2014a**).

In this study, we developed a new dCas9-based gene activation system by recruiting a CMV enhancer in *trans* to target gene via dCas9 associated with a simple chimeric sgRNA. This study revealed that the *trans* enhancer consisted of dCas9-VP64/csgRNA and sCMV could highly activate the expression of variant exogenous and endogenous genes in various mammalian cells, more efficiently than the currently widely used dCas9-VP64 and dCas9-VPR. This study thus developed a new strategy to activating endogenous genes, trans enhancer. To our knowledge, this is the first report to activate gene expression with an enhancer DNA in trans form. This creative strategy has its advantages over the current dCas9-based gene activation systems.

In this study, only one csgRNA was always used to activate all target genes in various cells. This differs from the reported dCas9/sgRNA gene activation systems. Currently, multiple sgRNAs have been employed in almost all reported gene activations with various dCas9/sgRNA systems. In general, three or more sgRNAs targeting to an interested gene are often used (**Cheng *et al*., 2013; Maeder *et al*., 2013; Mali *et al*. 2013a; Perez-Pinera *et al*., 2013**). In many comparative assays with various numbers of sgRNAs, one sgRNA produced very low or undetectable expression. In comparison with the current multiple sgRNAs strategy, our approach greatly simplified the sgRNA selection, design, and preparation, after all, not all genes are appropriate to design multiple sgRNAs in their promoter region. In addition, expression of multiple sgRNAs are difficult because each sgRNA has to independently transcribed by a long U6 promoter.

It can be seen that our csgRNA is a very simple engineered sgRNA in comparison with the currently reported engineered sgRNAs. The current dCas9/sgRNA activator systems often used a long complexed chimeric sgRNAs that harbor multiple tandem aptamers of various RNA-binding proteins, such as SAM sgRNA (MS2)(**Konermann *et al*., 2015; Liao *et al*., 2017**), Casilio sgRNA (Pumilio/FBF) (**Cheng *et al*., 2016**), and scaffold RNAs (MCP, PCP, and Com) (**Zalatan *et al*., 2015**). Our csgRNA only harbors a 24-bp short sequence at the end of normal sgRNA. It should be noted that we originally designed three different csgRNAs that have different capture sequences. We found that all of them work in the trans enhancer system; however, one (csgRNA2) showed the optimal performance. It should be pointed out that the capture sequences were artificially designed so that they have no complementary sequences in human cells, which is important for their high-efficiency specific annealing with sCMV. This study also reveals that the sCMV can anneal with csgRNA in the nuclear of human cells. This interaction successfully recruits the trans enhancer DNA to the target genes by dCas9/csgRNA. Clearly this interaction is more simple than that between RNA aptamers and RNA-binding proteins used in the current sCas9 activation systems. This develops a new approach for biomolecule recruitment in cells, which may be useful for future application.

It is clear that the *trans* enhancer technique is helpful for *in vitro* application, such as *in vitro* cell reprogramming and gene activation for gain-of function. However, it has to say that the *trans* enhancer still faces difficulty in *in vivo* application. The *trans* enhancer used a linear CMV enhancer DNA fragment that has a single-stranded overhang complementary to the 3’ end of csgRNA. It is difficult to produce this kind of *trans* enhancer DNA in the *in vivo* cells unless transfecting the *in vitro* pre-prepared *trans* enhancer with nanoparticle gene carriers together with expression vectors of dCas9-VP64 and csgRNA. However, the current *trans* enhancer can’t be brought into the *in vivo* cells by the current most effective *in vivo* transgenic vector, virus, such as AAV, that has been approved by FDA to clinical application. Other new strategies should be conceived to address the problem for the *in vivo* application of *trans* enhancer. For example, fusing a DNA-binding domain to dCas9 that can bind *trans* enhancer with its binding sites.

In this study, we selected 10 endogenous genes to be activated by *trans* enhancer, including *HNF4α, E47, Ascl1, Ngn2, Oct4, Sox2, Nanog, TNFAIP3(A20), CASP9*, and *CSF3*. These genes are not randomly selected. Most of these genes are transcription factors, including *HNF4α, E47, Ascl1, Ngn2, Sox2, Oct4*, and *Nanog*. The combination of *Ascl1, Ngn2*, and *Sox2* has been reported to be used to directly reprogram fibroblasts into nerve cells (**Zhao *et al*. 2015**). The combination of *Oct4, Sox2*, and *Nanog* has been widely used to reprogram fibroblasts into iPS cells (**Takahashi *et al*. 2007; Takahashi and Yamanaka 2016; Yu *et al*. 2007**). The *HNF4α* and *E47* have been used to differentiate the liver and panaceas cancers into normal cells (**Kim *et al*. 2015; Yin *et al*. 2008**). *TNFAIP3(A20)* is a well-known natural NF-κB inhibitor (**Cooper *et al*. 1996**), having the potential to treat NF-κB-over activated diseases such as inflammation and cancers. *Caspase9* is a key gene making cell apoptosis (**Li *et al*. 2017**). *CSF3* codes granulocyte-colony stimulating factor (G-CSF), a glycoprotein that stimulates the bone marrow to produce granulocytes and stem cells and release them into bloodstream (**Cetean *et al*. 2015**), which is widely used in chemotherapy to enhancer the immunity of cancer patients. This study revealed that the genes including *HNF4α, *E47*, Ascl1, Ngn2, TNFAIP3(A20)*, and *CSF3* were highly activated by *trans* enhancer in all transfected cells. However, the pluripotency-related genes, including *Oct4, Sox2* and *Nanog*, and *CASP9* were not highly activated as others, which may be related to the tight control of these critical genes.

## Materials and methods

### Vector construction

To apply the CRISPR/dCas9 expression system to transfection in cells, plasmid containing sgRNA derived by U6 promoter was constructed. We cloned a lac operon sequence with BbsI and BsaI sites at both ends from empty pEASY-Blunt-simple (Transgen, CB101-01) using Pfu High fidelity polymerase (Transgen, AS221-01) with primers Lac-px-F and Lac-px-R (**Supplementary Table 1**). This lac operon sequence was ligated into px458 (Addgene plasmid ID: 42230) to construct px458-lac. The ligation product was transformed into DH5a and then screened the blue colonies. The px458-lac was verified by sequencing. Then we designed three flanking sequences. The flanking sequence was add to 3’-end of gRNA scaffold sequence by PCR amplification using a forward primer (U6-F) and one of three reverse primers containing different flanking sequences, U6-1-R, U6-2-R, and U6-3-R (**Supplementary Table 1**), in which the gRNA scaffold sequence cloned in the px458-lac was used as template. The products were cloned into the pEASY-Blunt-simple, which produced the csgRNA expression vector named as pEASY-csgRNA. The csgRNA refers to capture sgRNA that has a 3’ extended flanking sequence. All flanking sequences were 24-bp in length.

Ten vital genes, including *HNF4α, TCF3(E47), ASCL1, Ngn2, Oct4, Nanog, TNFAIP3, CASP9, CSF3*, and *Sox2*, were chosen as target. The sgRNA target sites specific to each gene was designed by CHOPCHOP (https://chopchop.rc.fas.harvard.edu/). The pEASY-csgRNA expressing a particular csgRNA was constructed with a procedure described by **Supplementary Figure 10**. The chemically synthesized complementary oligonucleotides (**Supplementary Table 2**) with a 20-bp target-specific region flanked by two BbsI sites were diluted to 10 μM and mixed at the same molar. The final concentration of each oligonucleotide in a reaction was 1 μM. After mixing well, the complementary oligonucleotides were denatured with high temperature (95 °C) and annealed through a natural cooling process. The hybridization product were diluted 800 times and then ligated into pEASY-csgRNA. The ligation reaction (10 μL) consisted of 1 μL BbsI, 0.3 μL T4 DNA ligase, 1 μL 10× T4 DNA ligase buffer, 10× Bovine Serum Albumen (final concentration of 0.1 mg/mL), 50 ng plasmid pEASY-csgRNA, and ddH2O up to 10 μl total reaction volume. The ligation reactions was run cycle as fellow: 10 rounds of 37°C 5 min and 16°C 10 min, 37°C 30 min, and 80°C 5 min. The ligation product was introduced into DH5α competent cells and white clones were screened by blue-white screening on LB agar plates with 100 μg/mL Ampicillin, 40 μL of 20 mg/mL X-gal and 40 μL of 0.1 M IPTG. The vectors were validated by sequencing. Then the linear sgRNA or csgRNA expression sequences were amplified from the plasmid pEASY-csgRNAs by PCR using a forward primer U6-F and one of reverse primer U6-R, U6-1-R, U6-2-R, and U6-3-R. The primer U6-F and U6-R were used to amplify the normal sgRNA expression template (named as U6-sgRNA), and the primer U6-F and U6-1/2/3-R were used to amplify the csgRNA expression template (named as U6-csgRNA). The PCR products were purified by PCR clean kit (Axygen). These sequences were used to transfect cells as sgRNA transcription templates.

The complete sequence of CMV enhancer/promoter was amplified from pEGFP-N1 by using a forward primer (CMV-F; **Supplementary Table 1**) and one of reverse primers that had a special flanking sequence (CMV-1-R, CMV-2-R, or CMV-3-R; **Supplementary Table 1**). The amplified CMV promoter fragment harbors an Nt.BbvCI site at its one end. Then the amplified CMV promoter fragment was digested by Nt.BbvCI. The digested CMV promoter fragment was mixed with a complementary oligonucleotide (CS-1-R, CS-2-R, or CS-3-R; **Supplementary Table 1**) and then denatured at 85°C for 10 min. The denatured mixture was then naturally cooled to room temperature. The complementary oligonucleotide has the same sequence with the final 3’ single-strand stick end of CMV promoter fragment. The complementary oligonucleotide was used to remove the denatured oligo from CMV promoter fragment. The CMV promoter fragment was purified with PCR clean kit and used as linear stick-end CMV (sCMV) promoter fragment. The untreated CMV promoter fragment was used as a blunt-end CMV (bCMV) control.

A ZsGreen reporter construct was constructed by cloning an *HNF4α* promoter sequence into the upstream of ZsGreen gene. A 1000-bp *HNF4α* promoter sequence was amplified from the genomic DNA of HepG2 cells by PCR using primers HNF4α-P-F and HNF4α-P-R (**Supplementary Table 1**). Then the promoter sequence was ligated into the pEZX-ZsGreen, which produced an *HNF4α* promoter reporter construct named as pEZX-HP-ZsGreen. The vector pcDNA-dCas9-VP64 was purchased from Addgene (plasmid ID: 47107). The VP64 sequence in the pcDNA-dCas9-VP64 was deleted to construct pcDNA-dCas9. The VP64-p65-Rta (VPR) transcription-activating domain sequence was ligated into pcDNA-dCas9 to construct pcDNA-dCas9-VPR.

### DNA cutting with Cas9/csgRNA

A sgRNA targeting to the *HNF4α* promoter sequence was selected. The sgRNAs were prepared by an in vitro transcription using T7 RNA polymerase (M0251S, NEB). The sgRNA transcription template was prepared by PCR amplification of sgRNA-coding sequence cloned in the sgRNA expression plasmid (pEASY-csgRNA) with a forward primer, *HNF4α*-T7-F (**Supplementary Table 1**), which containing a T7 promoter sequence (TAATACGACTCACTATAG, transcription beginning with the 3’ G), and one of four reverse primers, U6-R, U6-1-R, U6-2-R, and U6-3-R (**Supplementary Table 1**). A normal sgRNA (*HNF4α*-sgRNA) and three csgRNAs (*HNF4α*-csgRNAs) were prepared. A 732-bp HNF4α promoter fragments was amplified from pEZX-HP-ZsGreen-A by PCR using primers HNF4α-sP-F and HNF4α-sP-R (**Supplementary Table 1**). The Cas9 digestion reaction (30 μL) consisted of 1×Cas9 Nuclease Reaction Buffer, 1 μM Cas9 Nuclease (NEB, M0386T), and 300 nM HNF4α-sgRNA or HNF4α-csgRNA. The Cas9 nuclease reaction was firstly incubated at 25 °C for 10 min. The Cas9 reaction was then added with 400 ng of purified 732-bp HNF4α promoter fragment and incubated at 37 °C for 15 min. Finally, Cas9 nuclease was inactivated at 65 °C for 10 min. The reaction was run with 1.5% agarose gel electrophoresis.

### Cell culture and transfection

293T, HepG2, A549, SKOV3, HT29, PANC-1, and HeLa were obtained from the cell resource center of Shanghai Institutes for Biological Sciences, Chinese Academy of Sciences and maintained in an incubator set at 37 °C and 5% CO_2_. Cells were cultured in the Dulbecco’s Modified Eagle Medium (DMEM) or Roswell Park Memorial Institute (RPMI) 1640 medium supplemented with 10% FBS, 100 units/mL penicillin, and 100 μg/mL streptomycin. Cells at >70% confluency in each well of 12-well plate were transfected with 800 ng total DNA, including 500 ng plasmid (pcDNA-dCas9, pcDNA-dCas9-VP64, or pcDNA-dCas9-VPR), 150 ng linear sgRNA expression template (U6-sgRNA or U6-csgRNA), and 150 ng linear CMV, by Lipofectamine^®^ 2000 (ThermoFisher Scientific) using the following protocol. For each transfection, cells were cultured with 600 μL of Opti-MEM (ThermoFisher Scientific) at 37 °C for 0.5 h when cell grew to a density of 4×10^5^/well. A stock solution of 100 μL of Opti-MEM, 800 ng of total DNA, 100 μL of Opti-MEM, and 4 μL of Lipofectamine 2000 was made for per transfection. The solution was then vortexed and incubated for 5 min at room temperature. Thereafter, the Opti-MEM/Lipofectamine solution was added to the individual aliquots of DNA stocked in 100 μL of Opti-MEM, vortexed, and incubated for 20 min at room temperature before being added to each well. After incubated cell with transfection solution for 4 h, the medium of each well was replaced with 800 μL of fresh DMEM or RPMI 1640 medium containing 10% FBS. The cells were further incubated at 37 °C and 5% CO_2_ for another 36 h. At the end of experiment, all cells were observed and photographed under a fluorescence microscope (Olympus) at 200 × magnification.

CMV linear fragment with sticky end (sCMV) or with blunt end (bCMV) were used as activating factor mixed with other ingredients. In activating the *HNF4α* promoter reporter construct, the 293T cells were co-transfected with the HNF4α-sgRNA, pcDNA-dCas9-VP64, and pEZX-HP-ZsGreen as the VP64 control group, and the 293T cells were co-transfected with the HNF4α-csgRNA, pcDNA-dCas9-VP64, CMV linear fragment, and pEZX-HP-ZsGreen as trans-CMV group. In activating endogenous genes, cells were co-transfected with the sgRNA and pcDNA-dCas9-VP64 as the VP64 control group, and cells were co-transfected with the csgRNA, pcDNA-dCas9-VP64, and CMV linear fragment as trans-CMV group.

### Flow cytometry

The fluorescence intensity of cells was quantified with flow cytometry (Calibur, BD, USA). Ten thousand cells were measured for each transfection. Flow cytometry data analysis and figure preparation were done using BD software.

### Quantitative PCR

Total RNA was extracted from transfected cell using TRIzol™ (Invitrogen). cDNA was synthesized from up to 3 μg total RNA in a volume of 20 μl consisting of 1× Hifair™ III SuperMix (YEASEN, 11137ES50). All mRNA levels were normalized using GADPH mRNA as a control. Transcription levels of differently treated cells were analyzed by quantitative PCR using ABI Step One Plus (Applied Biosystems) according to the manufacturer’s protocol. The quantitative PCR used the primers listed in **Supplementary Table 2 and 3**.

### Wound healing migration assays

Cells were seeded in 24-well plates (Costar, Cambridge, MA) and grown to confluence (overnight). Then the cells were transfected with dCas9-VP64/sgRNA or dCas9-VP64/csgRNA & sCMV to activate the target genes. The cells were transfected for 4 hs. Then the transfection medium was removed and cells were cultured with the fresh complete medium for 36 hs. Cell monolayers were wounded with a sterile pipette tip and then rinsed with PBS to remove cellular debris. The wounded monolayers were cultured in complete media for 48 hs and images by microscope.

### Transwell assays

Cells were transfected with or without *trans* CMV enhancer to activate the target genes. After transfected 36 hours, the cells were removed medium and washed three times with PBS. Then cells were trypsinized and resuspended with medium. The resuspended cells diluted to 1×10^5^ cells/mL. In 24-well plate, each well was added 1 mL medium, and then transwell chamber was added into the well. Finally, 2×10^4^ cells (200 μL) were added into each transwell. Then the 24-well plate was maintained in an incubator set at 37 °C and 5% CO_2_ for 48 hs. The cells invaded through the transwell chamber and adhered to the surface of the well were fixed and stained with acridine orange and imaged with fluorescence microscope. Cells in acridine orange-stained images were counted by ImageJ.

### Acridine orange staining

After transfection and culturing in fresh medium, each cell line was washed twice with PBS, then stained with 100 μg/mL acridine orange (AO) for 10 min at room temperature. Then cells were observed and photographed under a fluorescence microscope (Olympus) at 200 × magnification.

### Statistical analyses

Each cell transfection for detecting gene expression activation by trans-enhancer was performed in three biological replicates. In each biological replicate, at least three technical replicates were performed. In qPCR detection of gene expression, the mean RQ value of technical replicates was used as the RQ value of one biological replicate. The mean RQ value of three biological replicates were used to calculate the final mean and standard deviation (SD). Data were analyzed by Student’s t-test when comparing 2 groups. Data were shown as mean ± SD and differences were considered significant at *P* < 0.05.

## Acknowledgments

This work was supported by the grants from the National Natural Science Foundation of China (Grant 61571119).

## Author contributions

X.H.X. performed experiments, analyzed data and wrote the manuscript. D.Y.W., W.D. and J.W. assisted with the preparation of reagents, vector construction, data analysis and manuscript preparation. J.K.W. conceptualized the project, designed and supervised the research, and wrote the manuscript. J.K.W. provided financial support for the project.

## Additional files

Supplementary files The file contains three tables, ten figures, and vector sequences.

## Supplementary infromation

**Supplementary Tables** (Primer used in this study)

**Supplementary Table 1.**
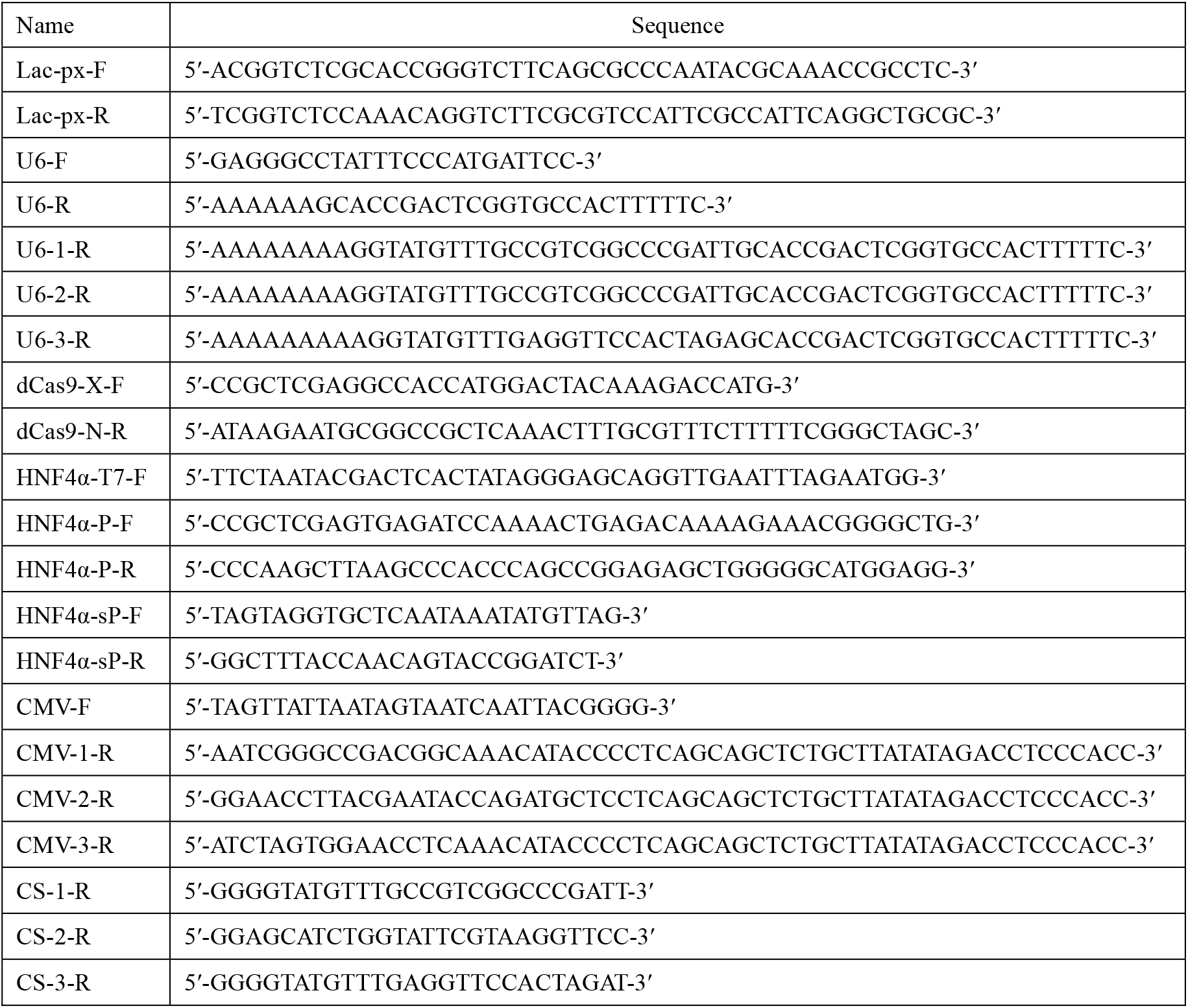
Primers were used in high fidelity amplification in this study.

**Supplementary Table 2.**
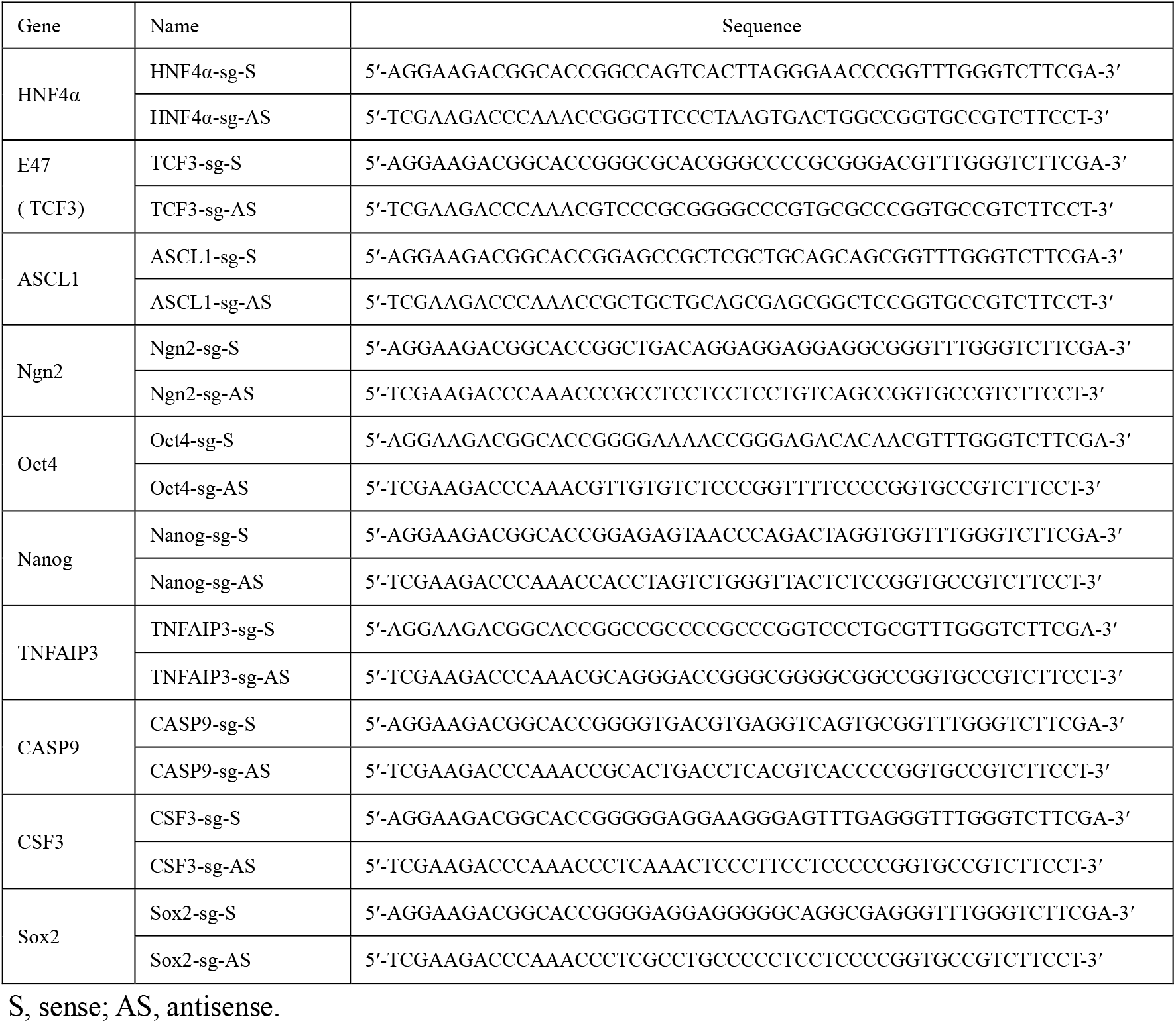
Oligonucleotide used to prepare target-specific regions (20 bp) of sgRNA

**Supplementary Table 3.**
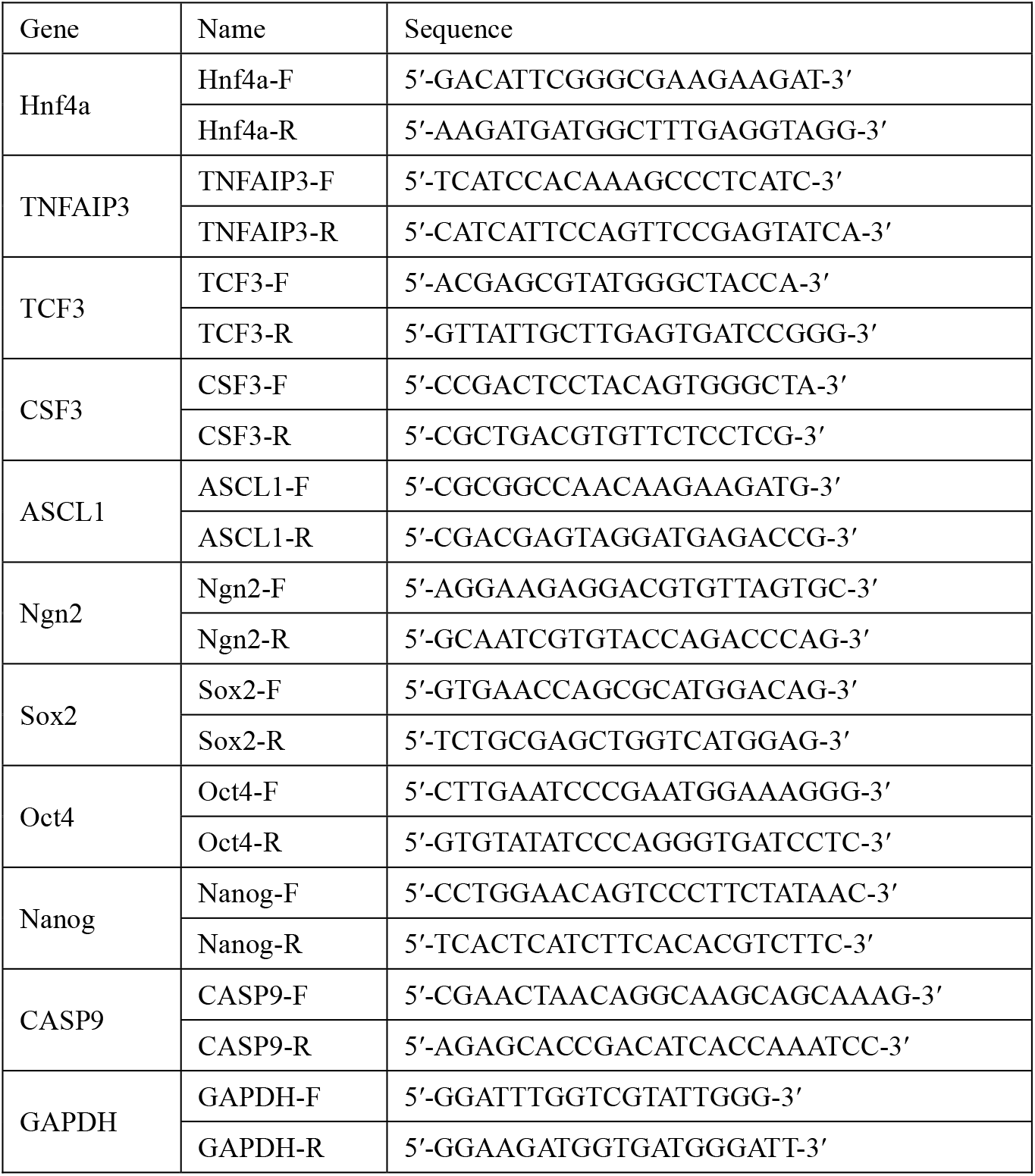
Primers used as qPCR detection of gene expression (see Figure 5).

**Supplementary Table 4.**
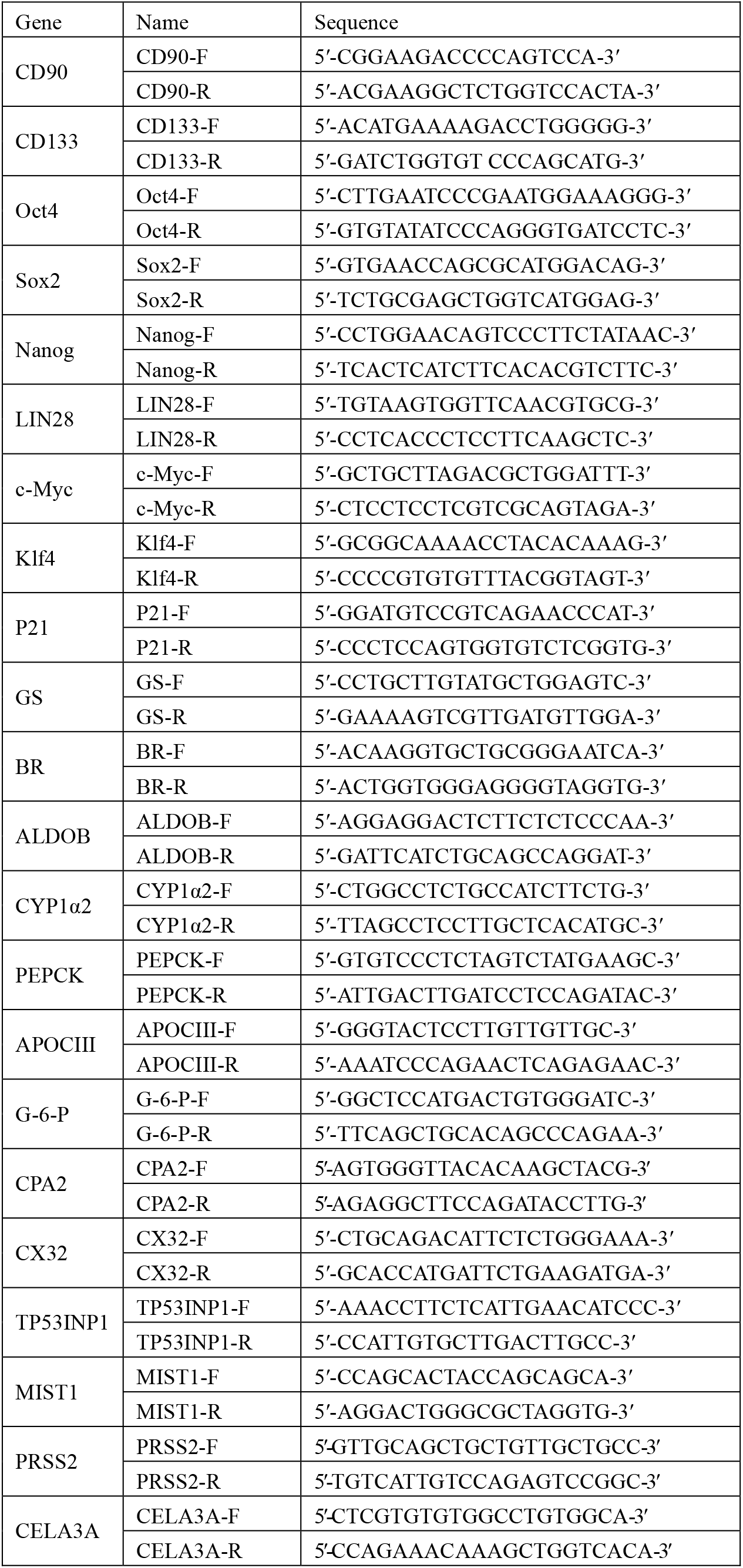
Primers used as qPCR detection of gene expression (see Figure 6).

## Supplementary Figures

**Supplementary Figure 1.**
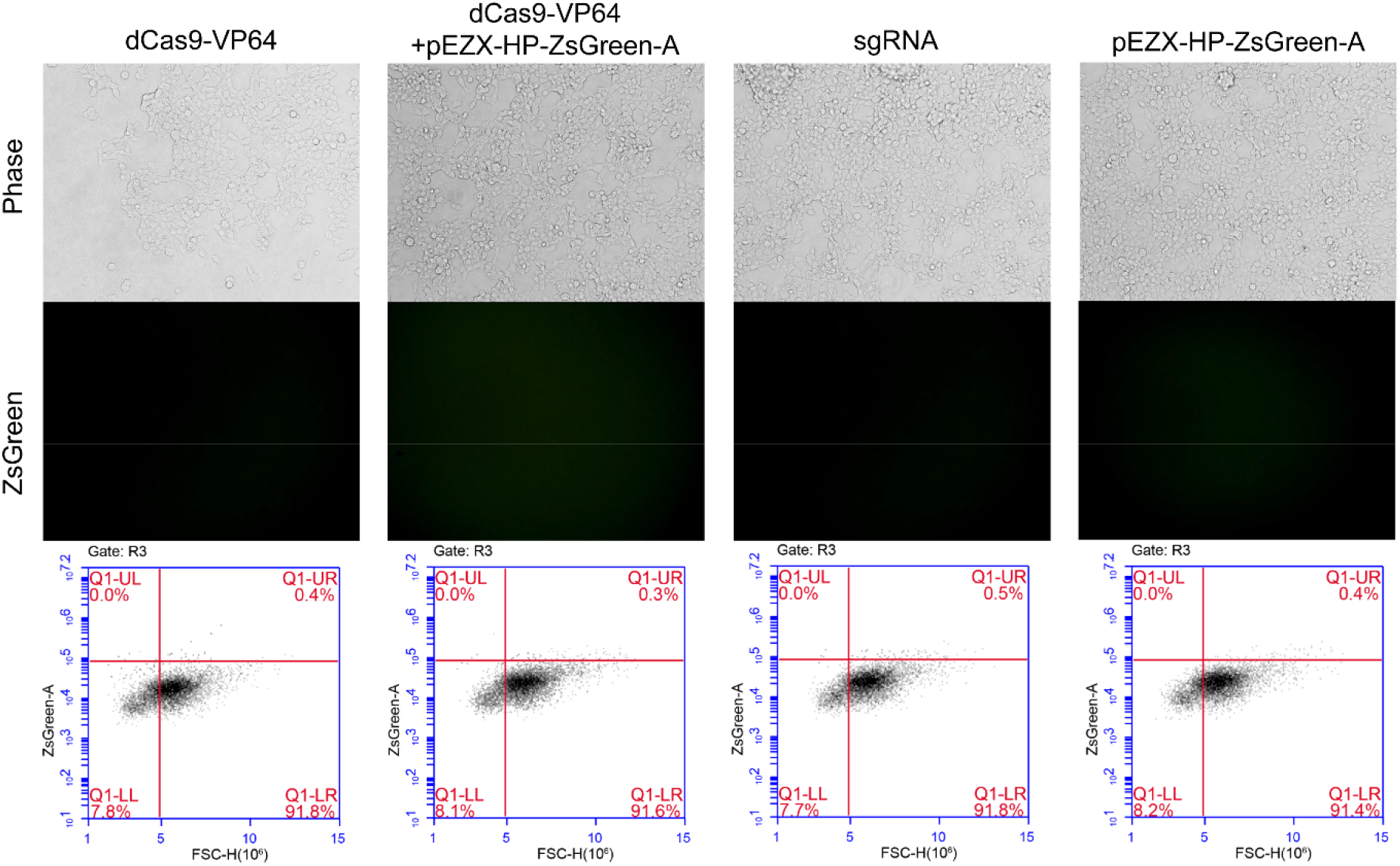
Activation of reporter gene ZsGreen under the control of *HNF4α* promoter in HepG2 cells with various activators. Cells were photographed and the fluorescence intensity was quantified by flow cytometry. The figure showed the images and fluorescence quantification results of cells as negative control transfections of Fig.3. The reporter gene activation efficiency was indicated by the percentage of cells with green fluorescence over the threshold (cells in Q1-UR quadrant).

**Supplementary Figure 2.**
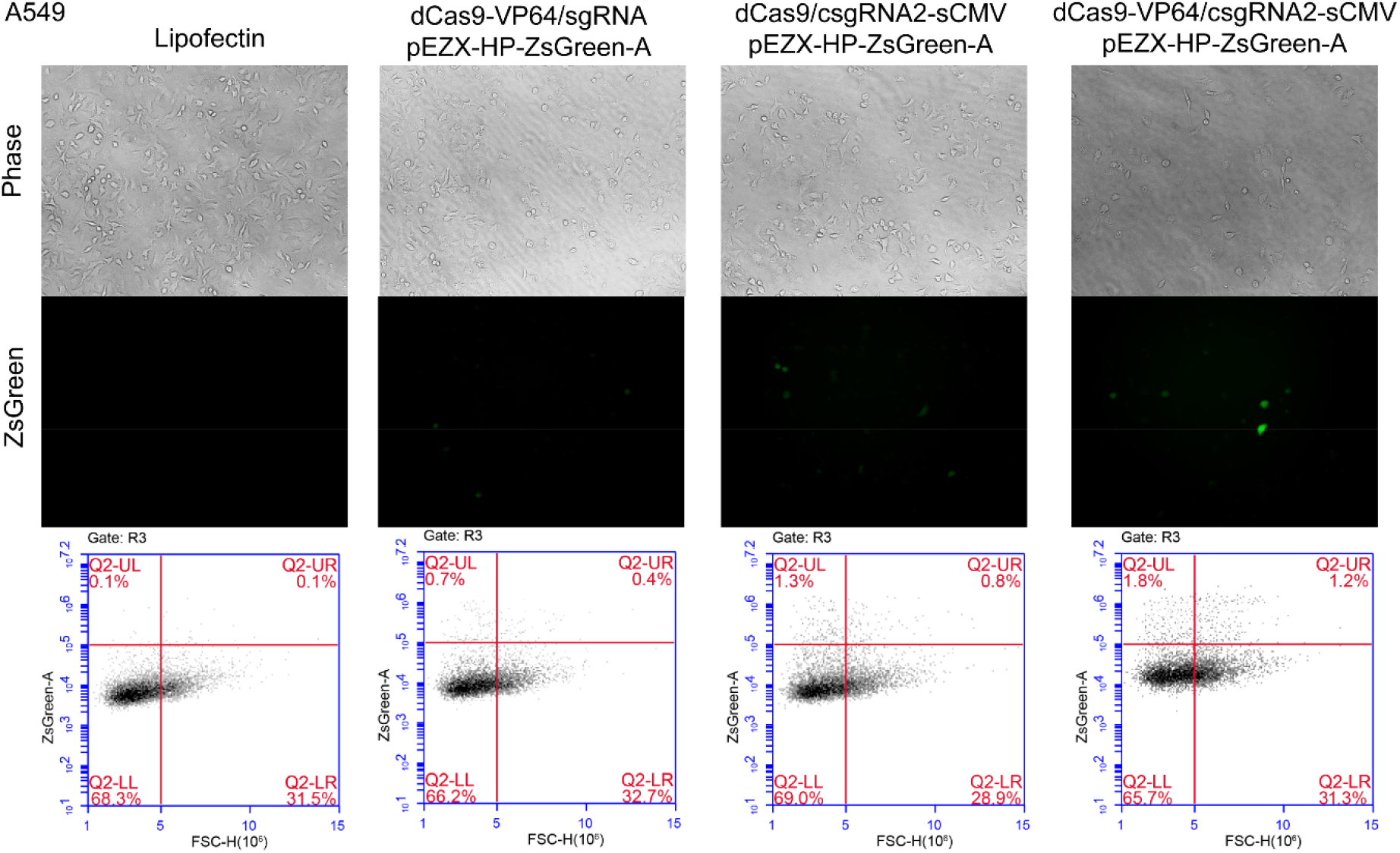
Activation of an exogenous reporter gene ZsGreen under the control of a *HNF4α* promoter by the CRISPR-assistant *trans* enhancer in the A549 cells. Cells were transfected with three transcriptional activation systems, including dCas9-VP64 & sgRNA, dCas9 & csgRNA & sCMV, and dCas9-VP64 & csgRNA2 & sCMV, together with a reporter construct (pEZX-HP-ZsGreen-A), respectively. Cells were photographed with a fluorescent microscope and their florescence intensity was analyzed by flow cytometry. The reporter gene activation efficiency was indicated by the percentage of cells with green fluorescence over the threshold (cells in Q1-UR quadrant).

**Supplementary Figure 3.**
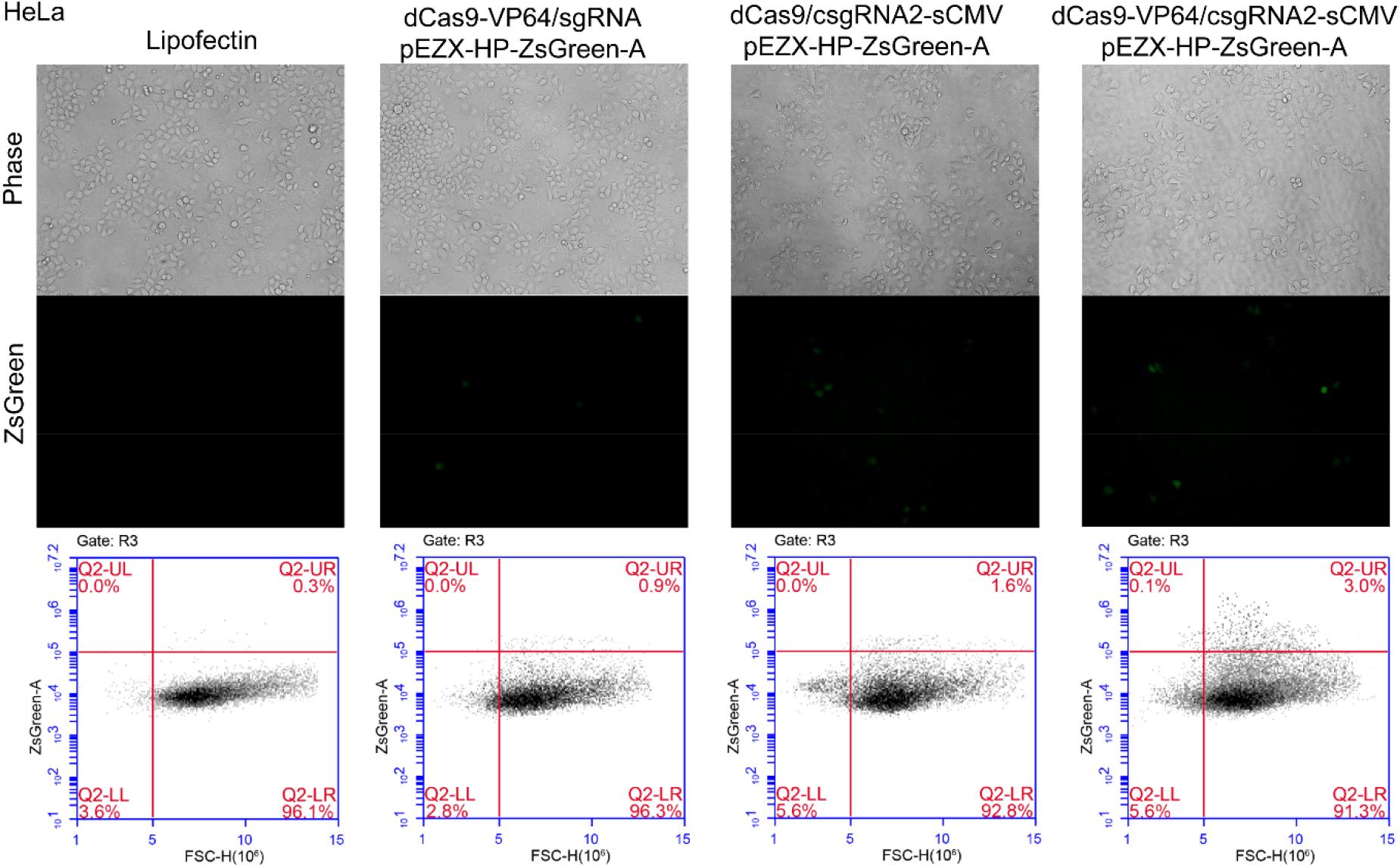
Activation of an exogenous reporter gene ZsGreen under the control of a *HNF4α* promoter by the CRISPR-assistant *trans* enhancer in the HeLa cells. Cells were transfected with three transcriptional activation systems, including dCas9-VP64 & sgRNA, dCas9 & csgRNA & sCMV, and dCas9-VP64 & csgRNA2 & sCMV, together with a reporter construct (pEZX-HP-ZsGreen-A), respectively. Cells were photographed with a fluorescent microscope and their florescence intensity was analyzed by flow cytometry. The reporter gene activation efficiency was indicated by the percentage of cells with green fluorescence over the threshold (cells in Q1-UR quadrant).

**Supplementary Figure 4.**
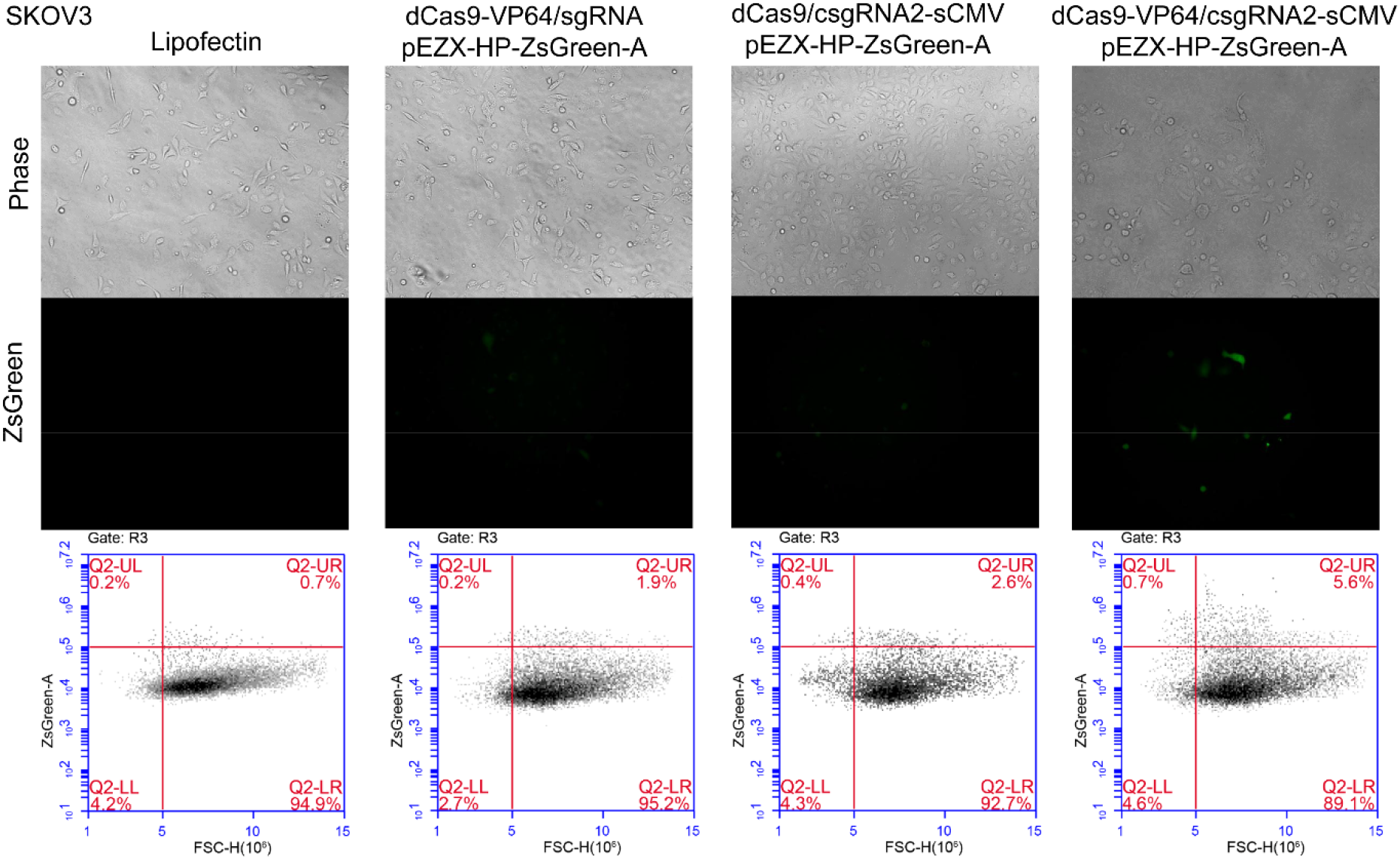
Activation of an exogenous reporter gene ZsGreen under the control of a *HNF4α* promoter by the CRISPR-assistant *trans* enhancer in the SKOV3 cells. Cells were transfected with three transcriptional activation systems, including dCas9-VP64 & sgRNA, dCas9 & csgRNA & sCMV, and dCas9-VP64 & csgRNA2 & sCMV, together with a reporter construct (pEZX-HP-ZsGreen-A), respectively. Cells were photographed with a fluorescent microscope and their florescence intensity was analyzed by flow cytometry. The reporter gene activation efficiency was indicated by the percentage of cells with green fluorescence over the threshold (cells in Q1-UR quadrant).

**Supplementary Figure 5.**
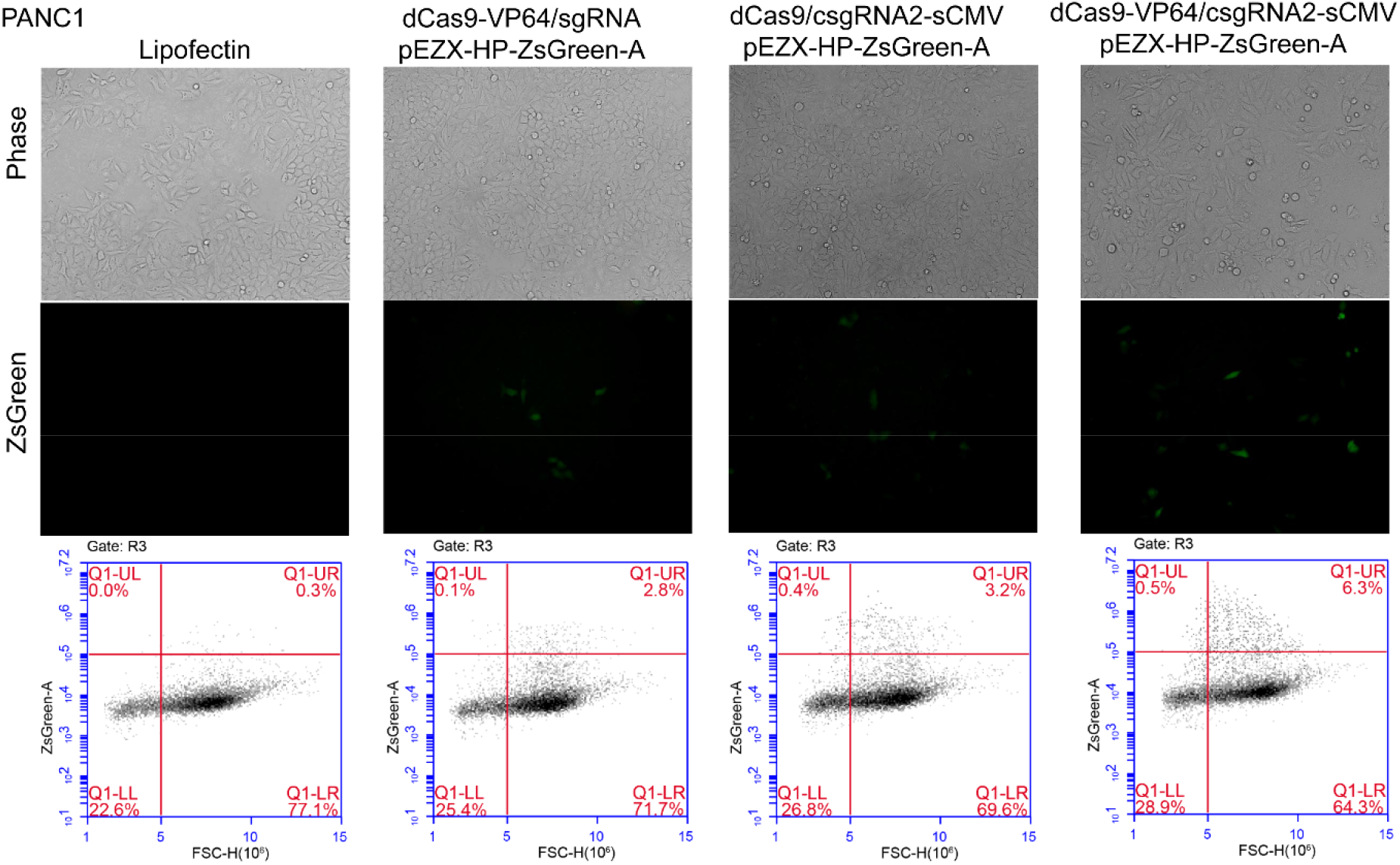
Activation of an exogenous reporter gene ZsGreen under the control of a *HNF4α* promoter by the CRISPR-assistant *trans* enhancer in the PANC1 cells. Cells were transfected with three transcriptional activation systems, including dCas9-VP64 & sgRNA, dCas9 & csgRNA & sCMV, and dCas9-VP64 & csgRNA2 & sCMV, together with a reporter construct (pEZX-HP-ZsGreen-A), respectively. Cells were photographed with a fluorescent microscope and their florescence intensity was analyzed by flow cytometry. The reporter gene activation efficiency was indicated by the percentage of cells with green fluorescence over the threshold (cells in Q1-UR quadrant).

**Supplementary Figure 6.**
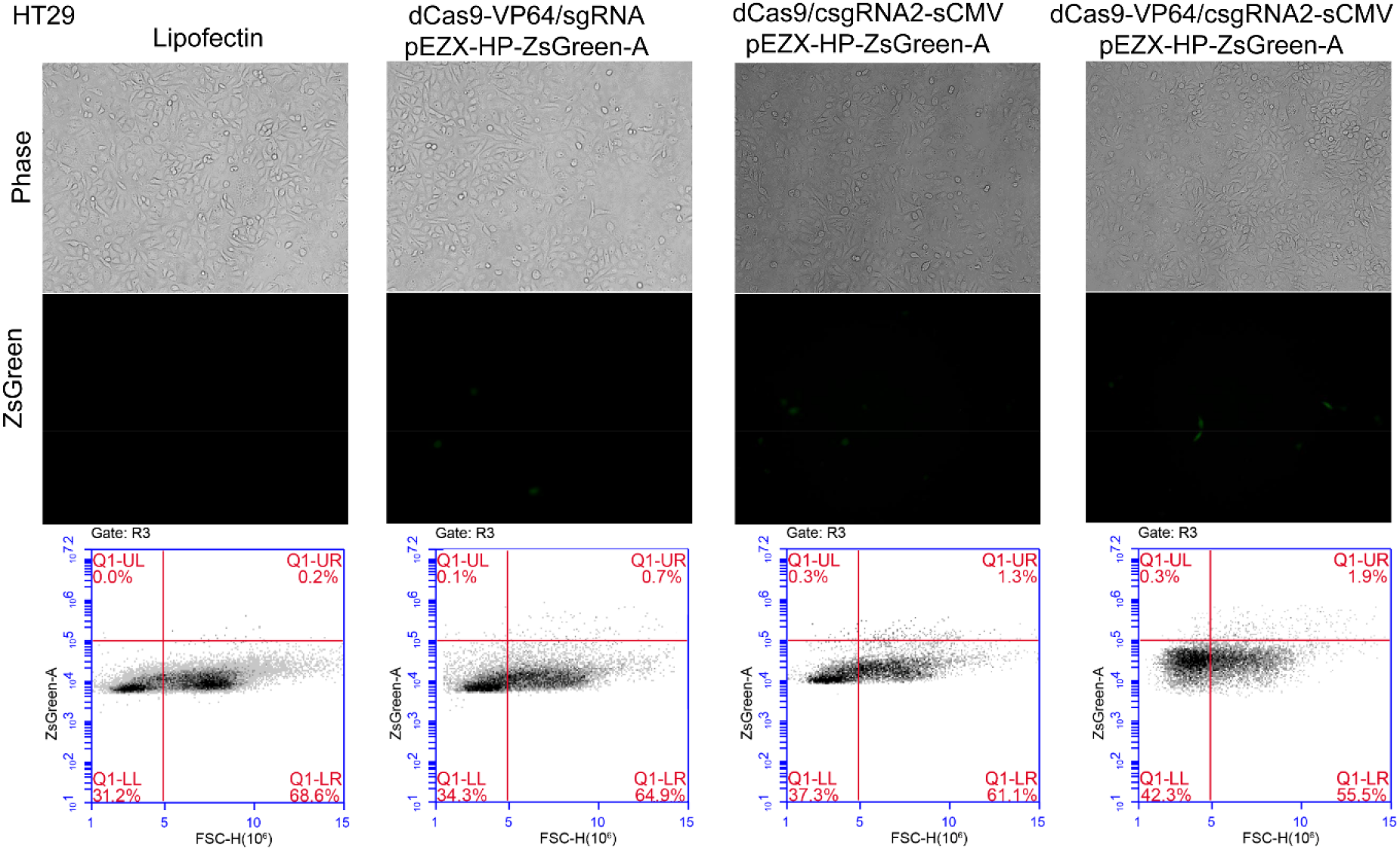
Activation of an exogenous reporter gene ZsGreen under the control of a *HNF4α* promoter by the CRISPR-assistant *trans* enhancer in the HT29 cells. Cells were transfected with three transcriptional activation systems, including dCas9-VP64 & sgRNA, dCas9 & csgRNA & sCMV, and dCas9-VP64 & csgRNA2 & sCMV, together with a reporter construct (pEZX-HP-ZsGreen-A), respectively. Cells were photographed with a fluorescent microscope and their florescence intensity was analyzed by flow cytometry. The reporter gene activation efficiency was indicated by the percentage of cells with green fluorescence over the threshold (cells in Q1-UR quadrant).

**Supplementary Figure 7.**
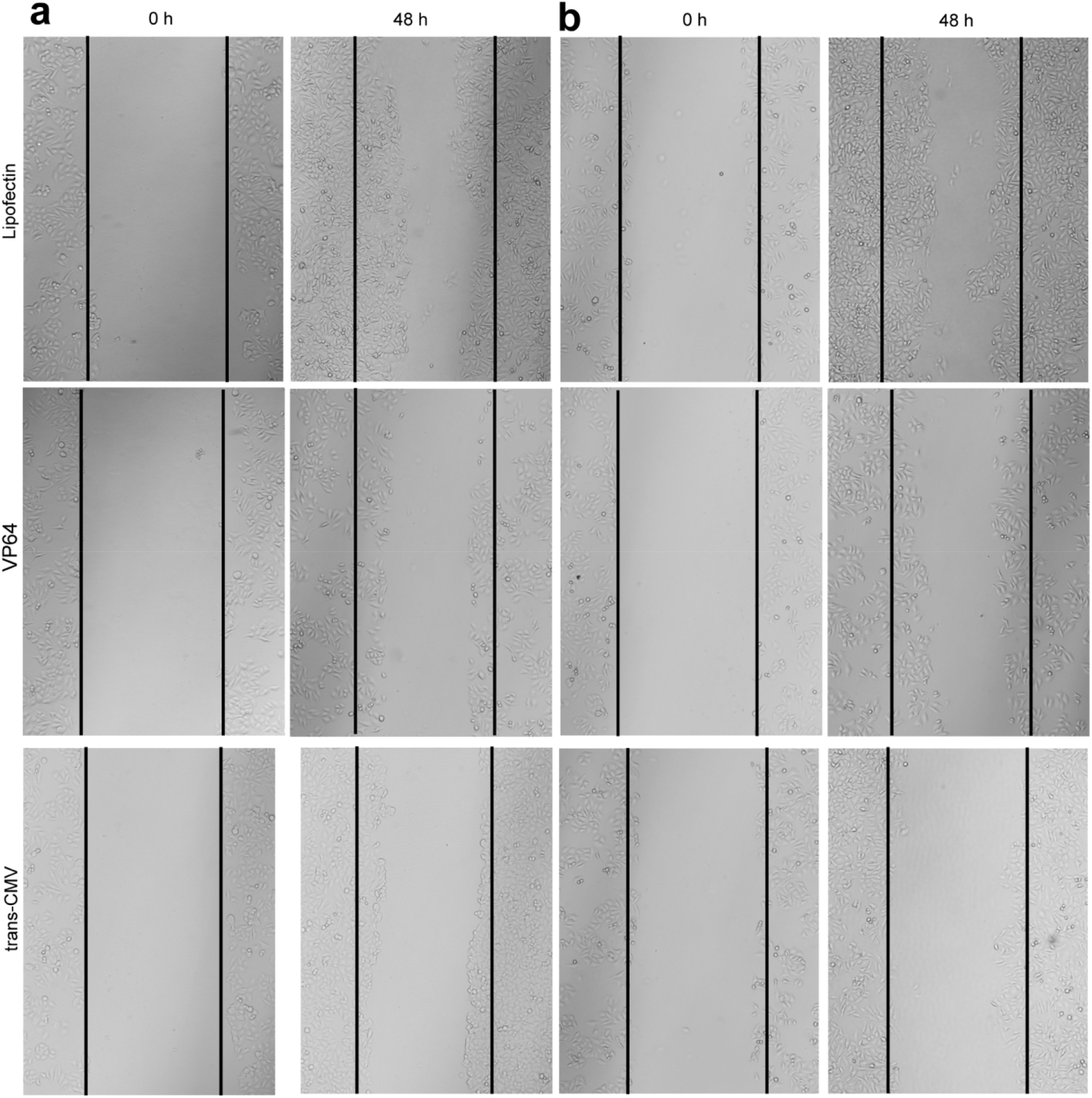
The wound-healing assay. Cells were transfected with dCas9-VP64/csgRNA (VP64) and dCas9-VP64/csgRNA&sCMV (Trans-CMV) to activate endogenous genes *HNF4α* in HepG2 cell and *E47* in in PANC-1 cell. a. The transferred HepG2 cells. b. The transferred PANC-1 cells.

**Supplementary Figure 8.**
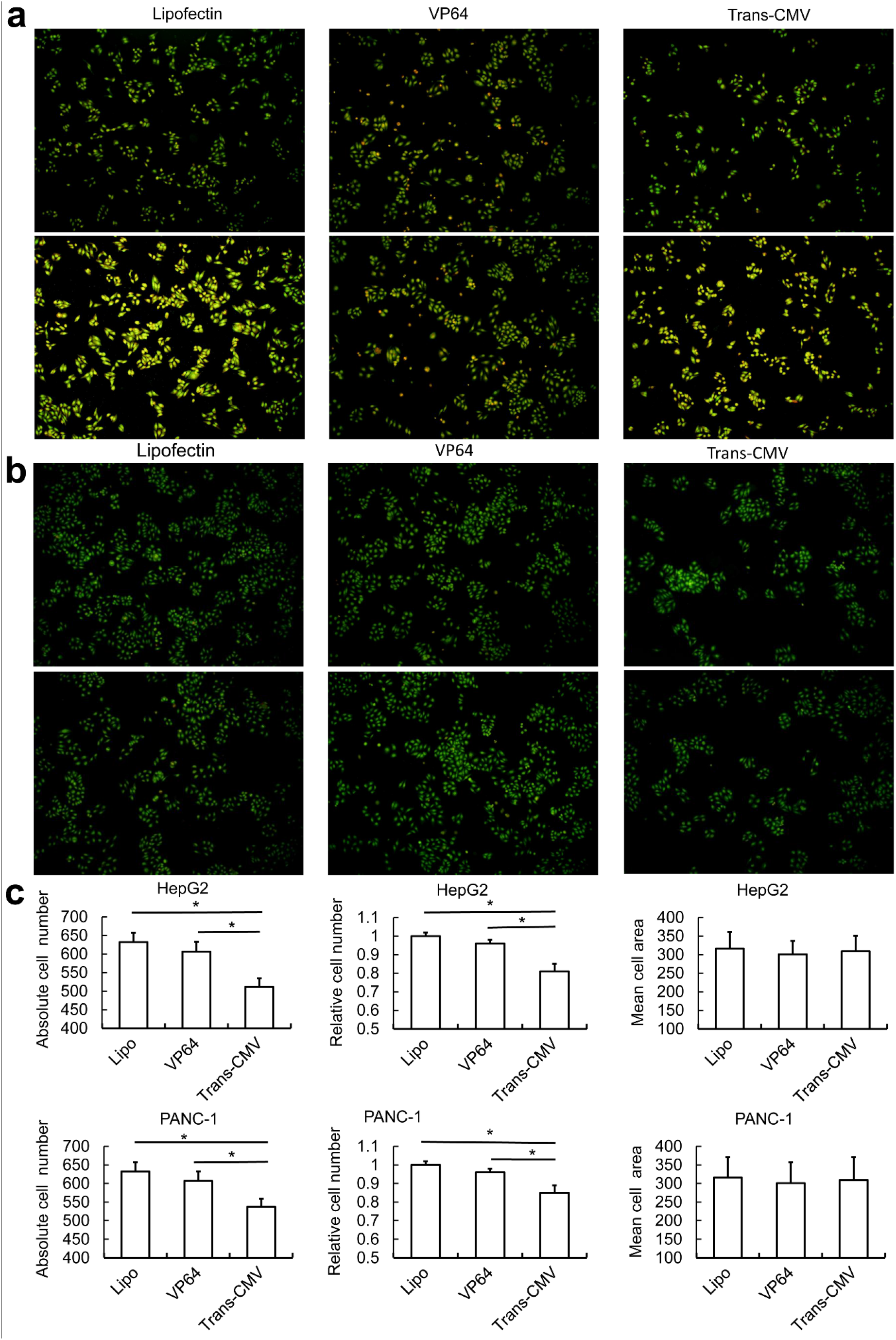
The transwell assay. Cells were transfected with dCas9-VP64/csgRNA (VP64) and dCas9-VP64/csgRNA&sCMV (Trans-CMV) to activate endogenous genes *HNF4α* in HepG2 cell and *E47* in in PANC-1 cell. a. The transferred HepG2 cells. b. The transferred PANC-1 cells. c. Counting of cell numbers. The images of acridine orange-stained cells were counted with ImageJ software. *, *p*<0.05.

**Supplementary Figure 9.**
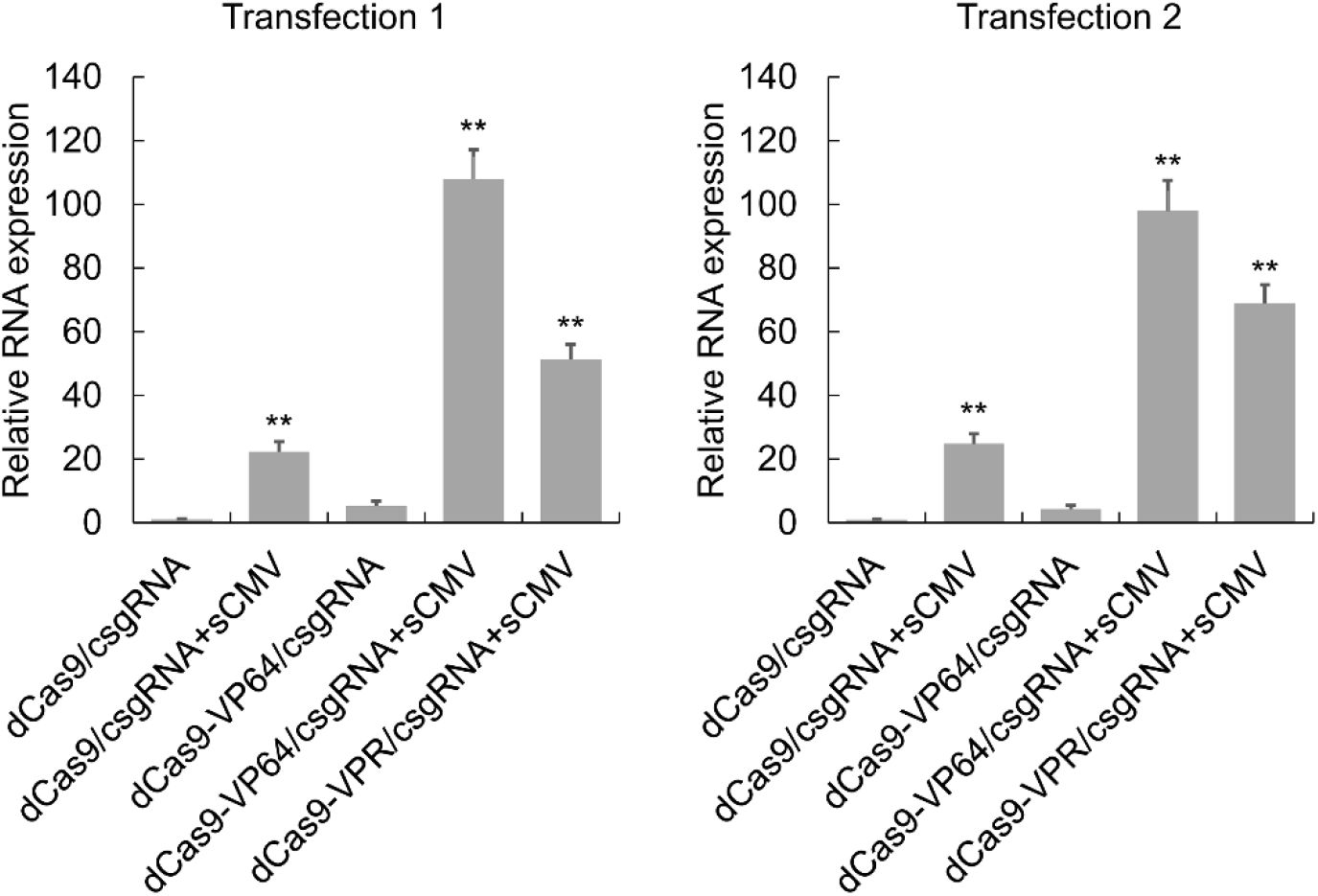
Activation of endogenous *HNF4α* gene in 293T cell with various systems. The 293T cell was transfected with various systems to activate the endogenous *HNF4α* gene. The *HNF4α* and β-action genes were detected with qPCR. The *HNF4α* expression level was showed as the fold of β-action.

**Supplementary Figure 10.**
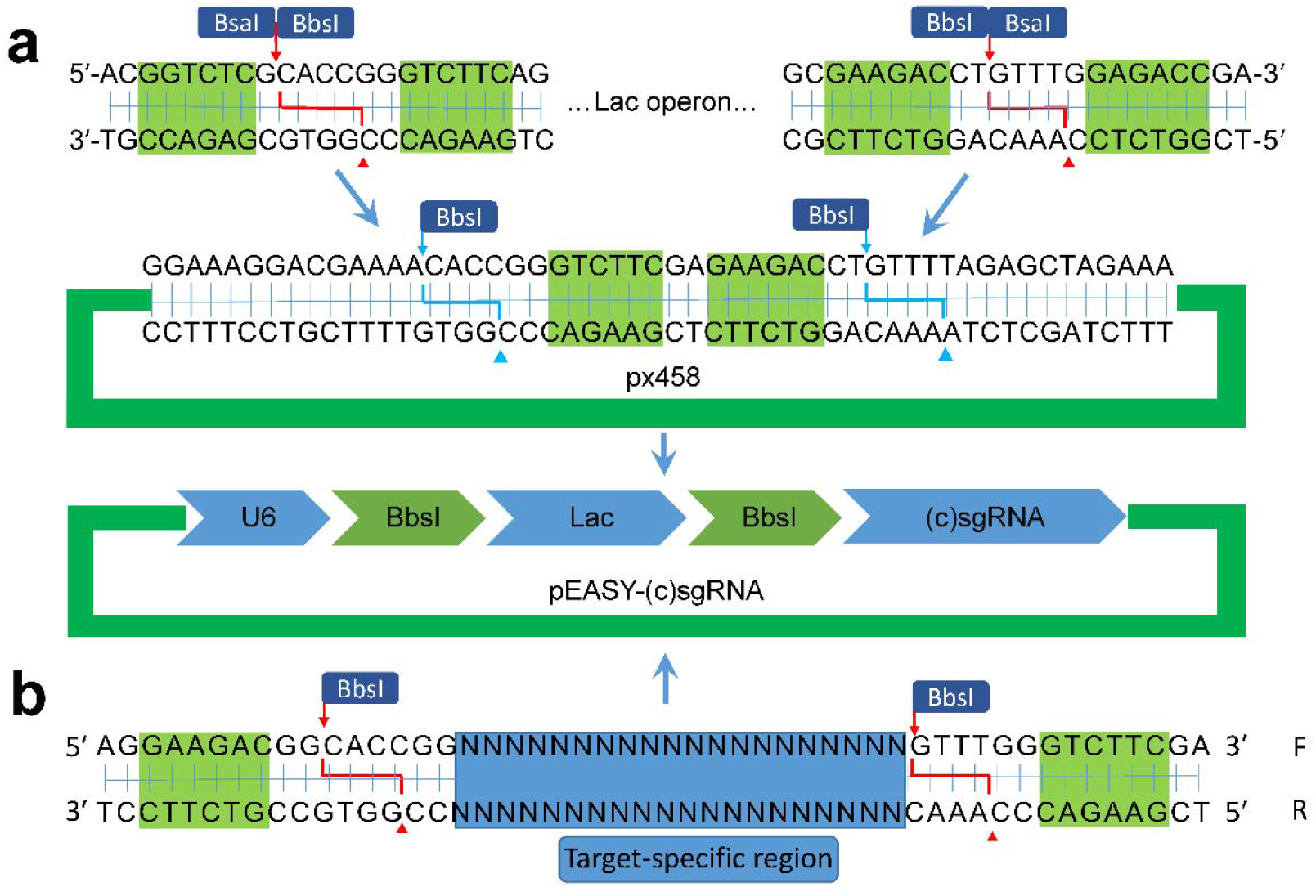
Construction of sgRNA vectors for blue-white screening.

## Supplementary sequences

### Sequences of vectors, templates, sgRNA, csgRNA, and sCMV

#### Receptor of pEASY-U6-csgRNA-1

U6 promoter + BbsI site + lac operon + BbsI site + gRNA scaffold + Flanking sequence 1 GAGGGCCTATTTCCCATGATTCCTTCATATTTGCATATACGATACAAGGCTGTTAGAGAGATAATTGGAATTAATTTGACTGTAAACACAAAGATATTAGTACAAAATACGTGACGTAGAAAGTAATAATTTCTTGGGTAGTTTGCAGTTTTAAAATTATGTTTTAAAATGGACTATCATATGCTTACCGTAACTTGAAAGTATTTCGATTTCTTGGCTTTATATATCTTGTGGAAAGGACGAAACACCGGGTCTTCA GCGCCCAATACGCAAACCGCCTCT CCCCGCGCGTTGGCCGATTCATTAAT GCAGCTGGCAC GACAGGTTTCCCGACTGGAAAGCGGGCAGTGAGCGCAACGCAATTAATGTGAGTTAGCTCACTCATTAGGCACCCCAGGCTTTACACTTTATGCTTCCGGCTCGTATGTTGTGTGGAATTGTGA GCGGATAACAATTTCACACAGGAAACAGCTATGACCATGATTACGCCAAGCTGCCCTTAAGG GCAGCTTCAATTCGCCCTATAGTGAGTCGTATTACAATTCACTGGCCGTCGTTTTACAACGTCGTGACTGGGAAAACCCTGGCGTTACCCAACTTAATCGCCTTGCAGCACATCCCCCTTTCGCCAGCTGGCGTAATAGCGAAGAGGCCCGCACCGATCGCCCTTCCCAACAGTTGCGCAGCCTGAATGGCGAATGGACGCGAAGACCTGTTTTAGAGCTAGAAATAGCAAGTTAAAATAAGGCTAGTC CGTTAT CAACTTGAAAAAGTGGCACCGAGTCGGTGCAATCGGGCCGACGGCAAACATACCTT TTTT

#### Receptor of pEASY-U6-csgRNA-2

U6 promoter + BbsI site + lac operon + BbsI site + gRNA scaffold + Flanking sequence 2 GAGGGCCTATTTCCCATGATTCCTTCATATTTGCATATACGATACAAGGCTGTTAGAGAGATAATTGGAATTAATTTGACTGTAAACACAAAGATATTAGTACAAAATACGTGACGTAGAAAGTAATAATTTCTTGGGTAGTTTGCAGTTTTAAAATTATGTTTTAAAATGGACTATCATATGCTTACCGTAACTTGAAAGTATTTCGATTTCTTGGCTTTATATATCTTGTGGAAAGGACGAAACACCGGGTCTTCA GCGCCCAATACGCAAACCGCCTCT CCCCGCGCGTTGGCCGATTCATTAAT GCAGCTGGCAC GACAGGTTTCCCGACTGGAAAGCGGGCAGTGAGCGCAACGCAATTAATGTGAGTTAGCTCACTCATTAGGCACCCCAGGCTTTACACTTTATGCTTCCGGCTCGTATGTTGTGTGGAATTGTGA GCGGATAACAATTTCACACAGGAAACAGCTATGACCATGATTACGCCAAGCTGCCCTTAAGG GCAGCTTCAATTCGCCCTATAGTGAGTCGTATTACAATTCACTGGCCGTCGTTTTACAACGTCGTGACTGGGAAAACCCTGGCGTTACCCAACTTAATCGCCTTGCAGCACATCCCCCTTTCGCCAGCTGGCGTAATAGCGAAGAGGCCCGCACCGATCGCCCTTCCCAACAGTTGCGCAGCCTGAATGGCGAATGGACGCGAAGACCTGTTTTAGAGCTAGAAATAGCAAGTTAAAATAAGGCTAGTC CGTTAT CAACTTGAAAAAGTGGCACCGAGTCGGTGCGGAACCTTACGAATACCAGATGCTTT TTTT

#### Receptor of pEASY-U6-csgRNA-3

U6 promoter + BbsI site + lac operon + BbsI site + gRNA scaffold + Flanking sequence 3 GAGGGCCTATTTCCCATGATTCCTTCATATTTGCATATACGATACAAGGCTGTTAGAGAGATAATTGGAATTAATTTGACTGTAAACACAAAGATATTAGTACAAAATACGTGACGTAGAAAGTAATAATTTCTTGGGTAGTTTGCAGTTTTAAAATTATGTTTTAAAATGGACTATCATATGCTTACCGTAACTTGAAAGTATTTCGATTTCTTGGCTTTATATATCTTGTGGAAAGGACGAAACACCGGGTCTTCA GCGCCCAATACGCAAACCGCCTCT CCCCGCGCGTTGGCCGATTCATTAAT GCAGCTGGCAC GACAGGTTTCCCGACT GGAAAGCGGGCAGTGAGCGCAACGCAATTAATGTGAGTTAGCTCA CTCATTAGGCACCCCAGGCTTTACACTTTATGCTTCCGGCTCGTAT GTTGTGTGGAATTGTGA GCGGATAACAATTTCACACAGGAAACAGCTAT GACCATGATTACGCCAAGCTGCCCTTAAGG GCAGCTTCAATTCGCCCTATAGTGAGTCGTATTACAATTCACTGGCCGT CGTTTTACAACGTC GTGACTGGGAAAACCCTGGCGTTACCCAACTTAATCGCCTTGCAGCACATCCCCCTTTCGCCAGCTGGCGTAATAGCGAAGAGGCCCGCACCGATCGCCCTTCCCAACAGTTGCGCAGCCTGAATGGCGAATGGACGCGAAGACCTGTTTTAGAGCTAGAAATAGCAAGTTAAAATAAGGCTAGTCCGTTATCAACTTGAAAAAGTGGCACCGAGTCGGTGCATCTAGTGGAACCTCAAACATACCTTT TTT

#### Transcription template of normal sgRNA

U6 promoter + target sequence + gRNA scaffold GAGGGCCTATTTCCCATGATTCCTTCATATTTGCATATACGATACAAGGCTGTTAGAGAGATAATTGGAATTAATTTGACTGTAAACACAAAGATATTAGTACAAAATACGTGACGTAGAAAGTAATAATTTCTTGGGTAGTTTGCAGTTTTAAAATTATGTTTTAAAATGGACTATCATATGCTTACCGTAAC TTGAAAGTATTTCGATTTCTTGGCTTTATATATCTTGTGGAAAGGACGAAACACCGGN NNNNN NNNNNNNNNNNN NNGTTTTAGAGCTAGAAATAGCAAGTTAAAATAAGGCTAGTCCGTTATCAA CTTGAAAAAGTGGCACCGAGTCGGTGCTTTTTT

#### Transcription template of csgRNA-1

U6 promoter + target sequence + gRNA scaffold + Flanking sequence 1 GAGGGCCTATTTCCCATGATTCCTTCATATTTGCATATACGATACAAGGCTGTTAGAGAGATAATTGGAATTAATTTGACTGTAAACACAAAGATATTAGTACAAAATACGTGACGTAGAAAGTAATAATTTCTTGGGTAGTTTGCAGTTTTAAAATTATGTTTTAAAATGGACTATCATATGCTTACCGTAAC TTGAAAGTATTTCGATTTCTTGGCTTTATATATCTTGTGGAAAGGACGAAACACCGGN NNNNN NNNNNNNNNNNN NNGTTTTAGAGCTAGAAATAGCAAGTTAAAATAAGGCTAGTCCGTTATCAA CTTGAAAAAGTGGCACCGAGTCGGTGCAATCGGGCCGACGGCAAACATACCTTTTTT

#### Transcription template of csgRNA-2

U6 promoter + target sequence + gRNA scaffold + Flanking sequence 2 GAGGGCCTATTTCCCATGATTCCTTCATATTTGCATATACGATACAAGGCTGTTAGAGAGATAATTGGAATTAATTTGACTGTAAACACAAAGATATTAGTACAAAATACGTGACGTAGAAAGTAATAATTTCTTGGGTAGTTTGCAGTTTTAAAATTATGTTTTAAAATGGACTATCATATGCTTACCGTAAC TTGAAAGTATTTCGATTTCTTGGCTTTATATATCTTGTGGAAAGGACGAAACACCGGN NNNNN NNNNNNNNNNNN NNGTTTTAGAGCTAGAAATAGCAAGTTAAAATAAGGCTAGTCCGTTATCAA CTTGAAAAAGTGGCACCGAGTCGGTGCGGAACCTTACGAATACCAGATGCTTTTTTT

#### Transcription template of csgRNA-3

U6 promoter + target sequence + gRNA scaffold + Flanking sequence 3 GAGGGCCTATTTCCCATGATTCCTTCATATTTGCATATACGATACAAGGCTGTTAGAGAGATAATTGGAATTAATTTGACTGTAAACACAAAGATATTAGTACAAAATACGTGACGTAGAAAGTAATAATTTCTTGGGTAGTTTGCAGTTTTAAAATTATGTTTTAAAATGGACTATCATATGCTTACCGTAAC TTGAAAGTATTTCGATTTCTTGGCTTTATATATCTTGTGGAAAGGACGAAACACCGGN NNNN NNNNNNNNNNNNN NNGTTTTAGAGCTAGAAATAGCAAGTTAAAATAAGGCTAGTCCGTTATCAA CTTGAAAAAGTGGCACCGAGTCGGTGCAT CTAGTGGAACCTCAAACATACCTTTTTT

#### sgRNA

target sequence + gRNA scaffold NNNNNNNNNNNNNNNNNNNNGUUUUAGAGCUAGAAAUAGCAAGUUAAAAUAAGGCUAGUC CGUUAUCAACUUGAAAAAGUGGCACCGAGUCGGUGCUUUU

#### csgRNA-1

target sequence + gRNA scaffold + Capture sequence 1 NNNNNNNNNNNNNNNNNNNNGUUUUAGAGCUAGAAAUAGCAAGUUAAAAUAAGGCUAGUC CGUUAUCAACUUGAAAAAGUGGCACCGAGUCGGUGCAAUCGGGCCGACGGCAAACAUAC CUUUU

#### csgRNA-2

target sequence + gRNA scaffold + Capture sequence 2 NNNNNNNNNNNNNNNNNNNNGUUUUAGAGCUAGAAAUAGCAAGUUAAAAUAAGGCUAGUC CGUUAUCAACUUGAAAAAGUGGCACCGAGUCGGUGCGGAACCUUACGAAUACCAGAUGC UUUUU

#### csgRNA-3

target sequence + gRNA scaffold + Capture sequence 3 NNNNNNNNNNNNNNNNNNNNGUUUUAGAGCUAGAAAUAGCAAGUUAAAAUAAGGCUAGUC CGUUAUCAACUUGAAAAAGUGGCACCGAGUCGGUGCAUCUAGUGGAACCUCAAACAUAC CUUUU

#### sCMV-1

CMV enhancer + CMV promoter + Nt.BbvCI site + Flanking sequence 1 5’-TAGTTATTAATAGTAATCAATTACGGGGTCATTAGTTCATAGCCCATATATGGAGTTCCGCGT TACATAACTTACGGTAAATGGCCCGCCTGGCTGACCGCCCAACGACCCCCGCCCATTGACG TCAATAAT GACGTATGTTCCCATAGTAACGCCAATAGGGACTTTCCATTGACGTCAAT GGGTG GAGTATTTACGGTAAACTGCCCACTTGGCAGTACATCAAGTGTATCATATGCCAAGTACGCCC CCTATTGACGTCAAT GACGGTAAATGGCCCGCCTGGCATTATGCCCAGTACAT GACCTTATGG GACTTTCCTACTTGGCAGTACATCTACGTATTAGTCAT CGCTATTACCATGGTGATGCGGTTTT GGCAGTACATCAATGGGCGTGGATAGCGGTTTGACTCACGGGGATTTCCAAGTCTCCACCC CATTGACGTCAAT GGGAGTTTGTTTTGGCACCAAAATCAACGGGACTTT CCAAAAT GTCGTAA CAACTCCGCCCCATTGACGCAAATGGGCGGTAGGCGTGTACGGTGGGAGGTCTATATAAGC AGAGCTGCTGAGGGGTATGTTTGCCGTCGGCCCGATT

#### sCMV-2

CMV enhancer + CMV promoter + Nt.BbvCI site + Flanking sequence 2 TAGTTATTAATAGTAATCAATTACGGGGTCATTAGTTCATAGCCCATATATGGAGTTCCGCGTTA CATAACTTACGGTAAATGGCCCGCCTGGCTGACCGCCCAACGACCCCCGCCCATTGACGTC AATAAT GACGTATGTTCCCATAGTAACGCCAATAGGGACTTTCCATTGACGTCAAT GGGTGGA GTATTTACGGTAAACTGCCCACTTGGCAGTACATCAAGTGTATCATATGCCAAGTACGCCCCC TATTGACGTCAAT GACGGTAAATGGCCCGCCT GGCATTATGCCCAGTACAT GACCTTATGGGA CTTTCCTACTTGGCAGTACATCTACGTATTAGTCAT CGCTATTACCATGGTGATGCGGTTTTGG CAGTACATCAATGGGCGTGGATAGCGGTTTGACTCACGGGGATTTCCAAGTCTCCACCCCATTGACGTCAATGGGAGTTTGTTTTGGCACCAAAATCAACGGGACTTTCCAAAATGTCGTAACAACTCCGCCCCATTGACGCAAATGGGCGGTAGGCGTGTACGGTGGGAGGTCTATATAAGCAGAGCTGCTGAGGAGCATCTGGTATTCGTAAGGTTCC

#### sCMV-3

CMV enhancer + CMV promoter + Nt.BbvCI site + Flanking sequence 3 TAGTTATTAATAGTAATCAATTACGGGGTCATTAGTTCATAGCCCATATATGGAGTTCCGCGTTACATAACTTACGGTAAATGGCCCGCCTGGCTGACCGCCCAACGACCCCCGCCCATTGACGTC AATAAT GACGTATGTTCCCATAGTAACGCCAATAGGGACTTTCCATTGACGTCAAT GGGTGGA GTATTTACGGTAAACTGCCCACTTGGCAGTACATCAAGTGTATCATATGCCAAGTACGCCCCC TATTGACGTCAAT GACGGTAAATGGCCCGCCT GGCATTATGCCCAGTACAT GACCTTATGGGA CTTTCCTACTTGGCAGTACATCTACGTATTAGTCAT CGCTATTACCATGGTGATGCGGTTTTGG CAGTACATCAATGGGCGTGGATAGCGGTTTGACTCACGGGGATTTCCAAGTCTCCACCCCATTGACGTCAATGGGAGTTTGTTTTGGCACCAAAATCAACGGGACTTTCCAAAATGTCGTAACAACTCCGCCCCATTGACGCAAATGGGCGGTAGGCGTGTACGGTGGGAGGTCTATATAAGCAGAGCTGCTGAGGGGTATGTTTGAGGTTCCACTAGAT

#### HNF4a promoter report vector

HNF4a promoter + minimal TATA-box promoter + ZsGreen TGAGAT CCAAAACTGAGACAAAAGAAACGGGGCTGTTCCAAAAAAAAAGCTAGGTGGCAGG TGTCTAACATGCCAGGGAGCTAAAACAGAGTGTGTGAGTTTCAGCAGCAGGTTGAATTTAGAATGGGGAAGGAGACCAGAGGAGACGCCAGACAGGATGACTTTGTCCCATTGGCCTGGAGGCAGCCCCATGTTTCTCCACCCCTCATATCACTCACCAGTTTGTAATAGTATCTTTGAATGACGATCTGATTAAGGTCCGTCTCCTCCATTAGTCCACAAGTTTCGGGGGTACATCTACTTTGCTCAT TTCCATATCCCCAGAGTCTAGCACAAGGCCTGGTACATAGTAGGTGCTCAATAAATATGTTAGA TGAAAGGAAGATAACACCTCTATGTACTAGCAGTGAGACTCCAGGCATGCAATTTCTCTCTGT CCTTCAGT CCCTTCAT CTCAAGGTTTAATTTAAATATGGTAACGCCTGTAT GCAACTCCCAGCA TCCAGTAGGCACTCACTAAACACAGTTCTCCACCCTCCTTTTTTCCTCTGCCCCTCCCTCGGTTTTCCCACTACTTCCTGCATGGTGACACACCCATAGTTTGGAGCCATAAAACCCAACCCAGGTTGGACTCTCACCTCTCCAGCCCCTTCTGCTCCGGCCCTGTCCTCAAATTGGGGGGCTGATGTCCCCATACACCTGGCTCTGGGTTCCCCTAACCCCAGAGTGCAGGACTAGGACCCGAGTGGACCTCAGGTCTGGCCAGGTCGCCATTGCCATGGAGACAGCAACAGTCCCCAGCCGCGG GTTCCCTAAGTGACTGGTTACTCTTTAACGTATCCACCCACCTTGGGTGATTAGAAGAATCAAT AAGATAACCGGGCGGTGGCAGCTGGCCGCACTCACCGCCTTCCTGGTGGACGGGCTCCTGGTGGCTGTGCTGCTGCTGTGAGCGGGCCCCTGCTCCTCCATGCCCCCAGCTCTCCGGCTG GGTGGGCTT AAGCTTCTCGACTTCCAGCTTGGCATAGAGGGTATATAATGGAAGCTCGACTT CCAGAT CCGGTACTGTTGGTAAAGCCACCGGATCCAGCCACCAT GGCCCAGTCCAAGCACG GCCTGACCAAGGAGATGACCATGAAGTACCGCATGGAGGGCTGCGTGGACGGCCACAAGTTCGTGATCACCGGCGAGGGCATCGGCTACCCCTTCAAGGGCAAGCAGGCCATCAACCTGTGCGTGGTGGAGGGCGGCCCCTTGCCCTTCGCCGAGGACATCTTGTCCGCCGCCTTCATGTA CGGCAACCGCGTGTTCACCGAGTACCCCCAGGACATCGTCGACTACTTCAAGAACTCCTGCCCCGCCGGCTACACCTGGGACCGCTCCTTCCTGTTCGAGGACGGCGCCGTGTGCATCTGCAACGCCGACATCACCGTGAGCGTGGAGGAGAACTGCATGTACCACGAGTCCAAGTTCTACGGCGTGAACTTCCCCGCCGACGGCCCCGTGATGAAGAAGATGACCGACAACTGGGAGCCCTCCTGCGAGAAGATCATCCCCGTGCCCAAGCAGGGCATCTTGAAGGGCGACGTGAGCATGTACCTGCTGCTGAAGGACGGTGGCCGCTTGCGCTGCCAGTTCGACACCGTGTACAAGGCCAAGTCCGTGCCCCGCAAGATGCCCGACTGGCACTTCATCCAGCACAAGCTGACCCGCGAGGACCGCAGCGACGCCAAGAACCAGAAGTGGCACCTGACCGAGCACGCCATCGCCTCCGGCTC CGCCTTGCCCGCCGCGCACCCGGGTTACTCTAGAGTCGGGGCGGCCGGCTAG

